# Non-Refoldability is Pervasive Across the *E. coli* Proteome

**DOI:** 10.1101/2020.08.28.273110

**Authors:** Philip To, Briana Whitehead, Haley E. Tarbox, Stephen D. Fried

## Abstract

The foundational paradigm of protein folding is that the primary sequence of a protein contains all the information needed for it to adopt a specific native structure^1,2^. This remarkable property is explained by Anfinsen’s thermodynamic hypothesis^3,4^, which asserts that because native states minimize the Gibbs free energy of a protein molecule under physiological conditions, proteins can reliably navigate to their native structures and remain in those states by ergodically sampling their free energy landscapes^5^. Most experiments of protein folding – conducted on purified, small, single-domain soluble proteins – follow the proportion of protein molecules that are folded as a function of time, temperature, denaturant concentration, or sequence^6–8^, and have yielded immense insight into the molecular determinants that underpin stable globular folds^9,10^. However, our reliance on the thermodynamic hypothesis as a ground truth to interpret these experiments has limited our ability to study the folding of complex proteins or to consider alternative non-thermodynamic scenarios^11,12^. Here, we introduce an experimental approach to probe protein refolding for whole proteomes. We accomplish this by first unfolding and refolding *E. coli* lysates, and then interrogating the resulting protein structures using a permissive protease that preferentially cleaves at flexible regions. Using mass spectrometry, we analyze the digestion patterns to globally assess structural differences between native and ‘refolded’ proteins. These studies reveal that following denaturation, many proteins are incapable of navigating back to their native structures. Our results signal a pervasive role for co-translational folding in shaping protein biogenesis, and suggest that the apparent stability of many native states derive from *kinetic persistence* rather than thermodynamic stability.

## Main Text

In our experiments (Fig. 1), *E. coli* cells are lysed under native conditions and the clarified extract divided so that one portion is retained in its native state, and a separate portion is first unfolded by addition of high concentrations of chemical denaturants (8 M urea or 6 M guanidinium chloride (GdmCl)) and then refolded by lowering the denaturant concentration (by dilution or by dialysis). Following a period of time, the structures of the proteins in these complex mixtures are probed by subjecting them to pulse proteolysis with proteinase K (PK). A low level of protease, active for a brief period of time (1 min), ensures that PK cleaves only at exposed or unstructured sites of target proteins, thereby encoding structural information of the protein’s conformation into cleavage sites^13^. After quenching PK, proteins are then fully trypsinized, and prepared for analysis by liquid chromatography/tandem mass spectrometry (LC-MS/MS). Half-tryptic peptides (HTPs), in which one cut-site is deemed to arise from trypsin (which only cuts after Arg and Lys) and the other cut-site from PK (which can cut between any two residues), reveal a location where PK cleaved. By quantifying the relative abundance of HTPs arising from the refolded sample compared to those from the native sample, one can assess sites in a protein where the local conformation is different in the refolded form. This method, called limited proteolysis mass spectrometry (LiP-MS) has been instrumental in exploring conformational change^14^, thermostability^15^, and allosteric binding^16^ on the proteome-scale, and here we have adapted it to probe protein folding.

**Fig. 1.**
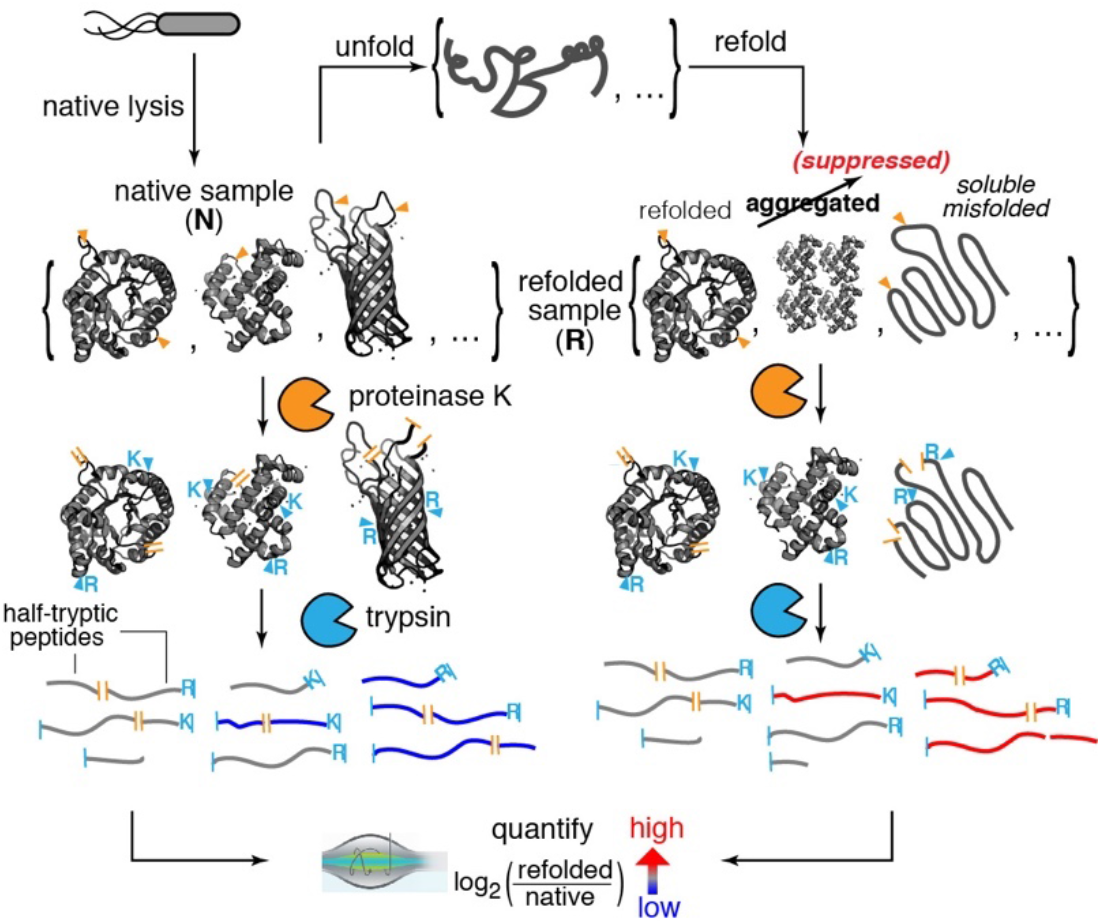
Limited Proteolysis Mass Spectrometry (LiP-MS) to Interrogate Refolding of the Proteome. Lysates from *E. coli* are prepared under native conditions, globally unfolded and refolded. The structures of the refolded proteins are probed by pulse proteolysis with proteinase K (PK) and compared to that of their native forms. Label free quantification (LFQ) of half-tryptic peptides reveals sites across the proteome where local conformation differ between a protein’s native and refolded forms.

A reversibly-refolding protein will have no ‘memory’ of being unfolded, and the refolded protein will equilibrate to the same ensemble of conformations. Therefore, the profile (and quantity) of HTPs will be identical between the native and refolded samples. On the other hand, aggregated proteins are expected to be more resistant to PK cleavage, whilst soluble misfolded proteins (with less compacted hydrophobic cores) are expected to be more susceptible to PK cleavage. To critically test these hypotheses, we first performed our LiP-MS method on two purified model proteins that are known refolders: Staphylococcal nuclease (SNase) and Ribonuclease H from *Thermus thermophilus* (*Tt*RNase H; Extended Data Fig. 1a). SNase refolded by dilution out of urea has a CD spectrum that superimposes on that of the native protein recombinantly expressed from *E. coli* (Fig. 2a). Likewise, when refolded and native SNase are probed with LiP-MS, they generate a set of 147 distinct tryptic and half-tryptic peptides (Fig. 2c) that are all present in equal abundances within our cut-offs for significance (abundance ratio greater than 2-fold, P-value by Welch’s t-test < 0.01). Importantly, when the same analysis is conducted on SNase spiked into *E. coli* lysate (Fig. 2d), the conformations of native and refolded protein are again indistinguishable. We repeated these studies on *Tt*RNase H, and found that its native and refolded forms generated overlapping CD spectra (Fig. 2b), and found no significant differences in their PK cleavage patterns – both in isolation (Fig. 2e; over 176 peptides) and when spiked into *E. coli* lysate (Fig. 2f; over 147 peptides). These studies show that LiP-MS provides a consistent picture with CD for refolding proteins, although does so with much greater structural resolution (providing independent quantifications at many distinct sites across the protein), and that thermodynamic refolding can be observed in a complex mixture^17^. Moreover, when these same experiments are conducted on SNase and *Tt*RNase refolded via slow dialysis from denaturant (rather than rapid dilution), we again found that the native and refolded forms were indistinguishable (Extended Data Fig. 1b-c), consistent with the notion that if a protein’s native state ensemble populates its free energy landscape’s thermodynamic minimum, correct refolding can occur regardless of the details of how the denaturant is removed.

**Fig. 2.**
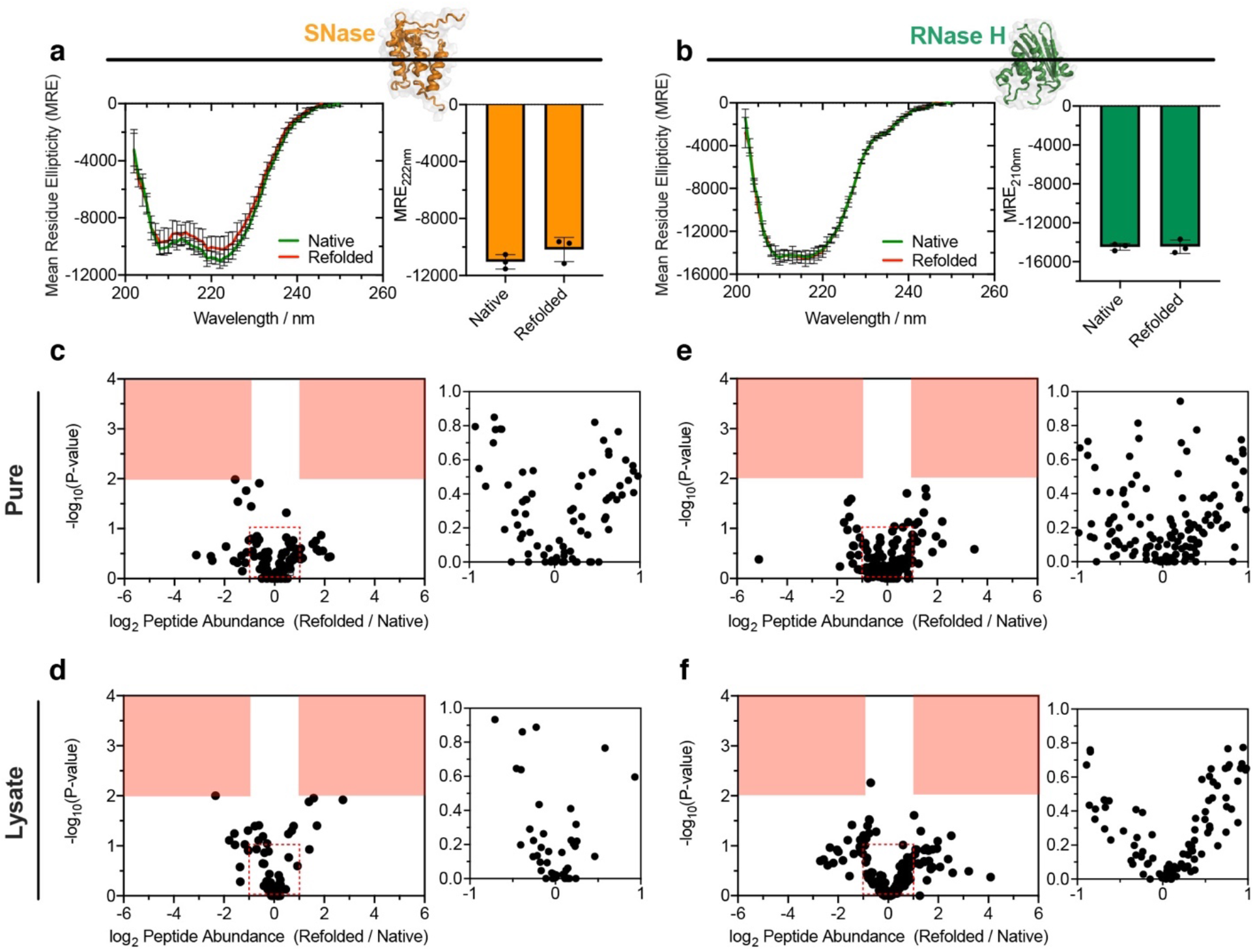
Refolding of Small Model Proteins. **a-b**, Circular Dichroism (CD) spectrum of **a**, Staphylococcal nuclease (SNase) or **b**, Ribonuclease H from *Thermus thermophilus* (*Tt*RNase H), natively expressed from *E. coli* (green), and following unfolding with 8 M urea and 50-fold dilution (red). Bar chart shows no significant difference in MRE at 222 nm (SNase) or 210 nm (RNase H) (*n* = 3). **c**, Volcano plot comparing peptide abundances from native and refolded SNase (*n* = 3). Effect sizes reported as ratio of averages, and P-values are based on Welch’s t-test. Red regions designate significance (effect-size > 2, P-value < 0.01). Inset shows large number of points clustered near the origin. The data suggest no significant difference in the structure of native SNase and the conformation produced when it is diluted out of urea. **d**, As in **c**, except SNase was spiked into *E. coli* lysate, providing a complex background. **e-f**, Volcano plot showing peptides quantified by LiP-MS derived from native and refolded *Tt*RNase H, both as a purified protein **e**, and spiked into *E. coli* lysate **f**.

### Refoldability of the Proteome

We next planned to carry out a similar experiment in which total soluble cellular extracts are unfolded, refolded, and then compared to a native lysate. However, a limitation that we considered is that many proteins in the *E. coli* lysate may aggregate during refolding. Proteins that aggregate could hypothetically interfere with other proteins in the mixture that could in principle refold on their own. To address this possibility, we unfolded and refolded *E. coli* lysates under a range of conditions and performed pelleting assays seeking conditions that minimized aggregation (Extended Data Fig. 2a-c). We found that precipitation could be minimized when refolding is conducted at an elevated pH (8.0) and low concentration (0.23 mg mL^−1^). We speculate this may be because the average isoelectric points (pI) among *E. coli* proteins is 6.6 ^18^, and so at pH 8.0 proteins will be preponderantly negatively charged, disfavoring nonspecific associations. Thermal denaturation followed by slow cooling generates substantially more precipitation (Extended Data Fig. 2c), suggesting such approaches may be better suited to probe aggregation propensity rather than individual proteins’ intrinsic refoldability^19^. Dynamic light scattering (DLS) experiments confirm that thermally-refolded lysates are highly enriched with large particle sizes (>0.2 *μ*m; Extended Data Fig. 2d), whilst chemically-refolded lysates had particle diameter distributions somewhat larger than (or similar to) native lysates. Collectively, these studies suggest refolding via removal of chemical denaturants at high dilution afford proteins in lysates an opportunity to refold in complex mixtures.

We performed proteome-wide studies (Fig. 1), unfolding *E. coli* lysates in both urea and GdmCl, initiating refolding by dilution, and recording refolding kinetics over time by pulsing samples with PK after discrete time increments. In these studies, we analyzed samples both with and without PK treatment. Comparison of the native and refolded samples without PK treatment enable the determination of overall changes in protein abundance (due to, for example, precipitation during refolding) and are used as normalization factors, so that abundance differences in peptide fragments can be attributed solely to structural changes in refolded proteins (Extended Data Fig. 2e,g). Consistent with our pelleting studies showing low levels of aggregation (Extended Data Fig. 2a), these normalization factors are almost all unity (Extended Data Fig. 2f,h). Because our conclusions rely heavily on missing data (a non-refolding protein will generate signature peptide fragments that are absent in its native form), we developed a new filtering algorithm for label free quantification that can reproducibly quantify peptide abundance over nine orders of magnitude (Extended Data Fig. 3a-d, Supplementary Note 1).

**Fig. 3.**
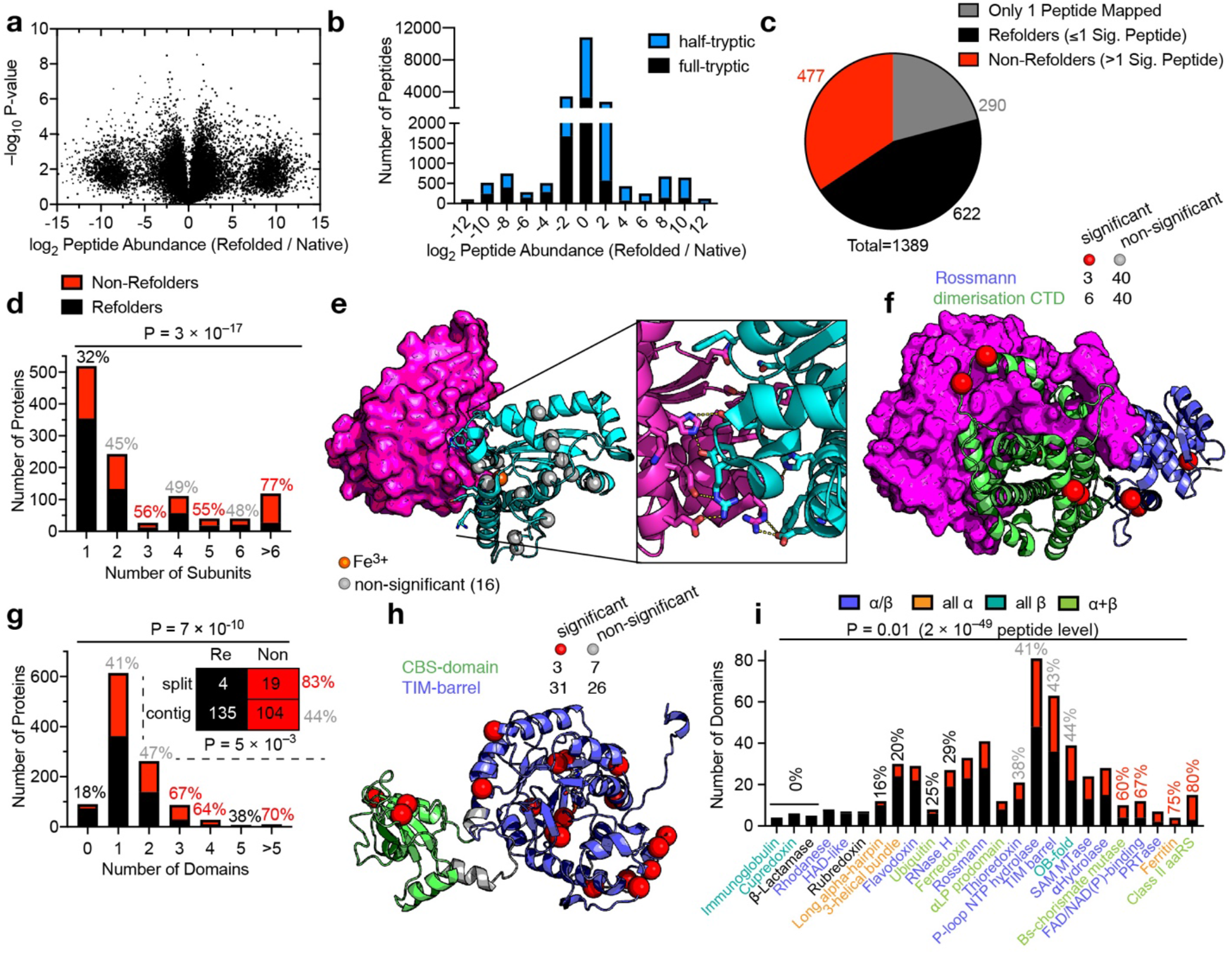
Refoldability of the *E. coli* Proteome. **a**, Volcano plot comparing peptide abundances from native and refolded *E. coli* lysates normalized for protein abundance (*n* = 3). Effect sizes reported as ratio of averages, and P-values are based on Welch’s t-test. **b**, Histogram of abundance ratios for half-tryptic and full-tryptic peptides. Half-tryptic peptides are enriched in the refolded proteome. **c**, Overall number of refolding proteins out of 1099 *E. coli* proteins. 290 proteins only furnished one peptide and hence too little data to make an assessment. **d**, Proteins that are part of complexes with multiple subunits – but especially trimers and pentamers – are significantly less refoldable than monomeric proteins (P = 3×10^−17^ by chi-square test). Percentages denote percent of proteins that are non-refoldable (red, higher than average; black, lower than average; gray, average). **e**, X-ray structure of Fe-superoxide dismutase (SodB, PDB: 2NYB), a refoldable homodimeric protein, showing a simple interface mediated primarily by polar interactions. **f**, X-ray structure of 6-phosphogluconate dehydrogenase (Gnd, PDB: 2ZYA), a non-refoldable homodimeric protein in which C-terminal domains (green) interdigitate. **g**, Proteins with many domains, but particularly three or four, are significantly more non-refoldable than proteins with zero or one domain (P = 7×10^−10^ by chi-square test). *Inset:* contingency table for 2-domain proteins comparing the refoldability of proteins in which domains are contiguous or in which one is split by an intervening domain. Split domains are strikingly less refoldable. **h**, X-ray structure of inosine 5’-monophosphate dehydrogenase (GuaB, PDB: 1ZFJ), a protein in which a split TIM-barrel domain is non-refoldable. **i**, Certain fold-types (e.g., ribonuclease H-like and immunoglobulin domains) are much more refoldable than others (e.g., ferritin-like and class II aaRS/biotin synthase-like). P = 0.01 by chi-square test though counts for many fold-types are low. Statistical significance is drastically enhanced by conducting analysis at the peptide level (see Extended Data Fig. 6).

Inspection of the dataset in which lysates were diluted out of GdmCl reveals that the distribution of abundance ratios for the 21,386 peptides quantified (Fig. 3a-b) follows a trimodal distribution, with half (50.6%) of the peptides present at similar abundances in the native and refolded samples (|log_2_ (Refolded / Native)| < 1), a sizable portion (7.8%) only detected in native samples, and another sizable portion (8.0%) only detected in the refolded samples. This distribution implies that many regions of *E. coli* proteins *failed* to refold, generating signature fragments that are completely absent in the native sample. HTPs (signifying exposed sites) are asymmetrically distributed and much more abundant in the refolded sample (Fig. 3b; P < 10^−218^ by chi-square test). This finding is consistent with the view that most proteins that fail to refold under our conditions form soluble disordered entities (rather than aggregate), which are more susceptible to PK cleavage.

We define a protein as ‘non-refoldable’ if we can map two or more peptides to it with significantly different abundances in the refolded sample, though our primary findings are not sensitive to this cut-off (Supplementary Data). Proteins with only one peptide mapped to it are considered to have too little data to make an assessment. Of the 21,386 peptides (mapping to 1,426 proteins) we quantified, 20,274 peptides (from 1,389 proteins) passed our stringent filters and were used for analysis. Among the set of 1,099 proteins with two or more mapped peptides, 477 (43%) were found not to be refoldable (Fig. 3c). Between technical replicates, ~90% of proteins were assigned the same refoldability status (Extended Data Fig. 3e-h).

We next sought to identify features of proteins that correlate with non-refoldability. Multimeric proteins showed a much lower proficiency to refold, whilst monomeric proteins had a high propensity to refold (P < 10^−16^ by chi-square test; Fig. 3d). Since natural protein folding on a polysome provides a means to couple folding and assembly, operon structure could facilitate biogenesis of multimeric proteins in a manner not recapitulated by *in vitro* refolding^20,21^. Dimers with simple or polar interfaces such as superoxide dismutase refold more easily (Fig. 3e) than dimers with complex interfaces (Fig. 3f) for which synthesis, folding, and assembly may require more coordination^22,23^. We found that trimers are much less refoldable than dimers and remarkably, also less refoldable than tetramers or hexamers. This may be because higher symmetry allows tetramers (or hexamers) to form in multiple discrete steps through stable intermediates^24^.

We also observed that proteins with more domains tend to refold significantly less efficiently (Fig. 3g), consistent with a prevailing view that the vectorial synthesis of translation decouples domains into independently folding units^25–27^, thereby preventing non-native inter-domain interactions that could form during refolding from denaturant. Strikingly, proteins with split domain architecture such as inosine 5’-monophosphate dehydrogenase (Fig. 3h) refolded exceptionally poorly (83% non-refoldable), suggesting a scenario where co-translational coordination is most critical (Extended Data Fig. 4a). Moreover, we observe evidence for coupling of domain refolding, such that domains in a multidomain protein more frequently all refold or all fail to refold (Extended Data Fig. 4b-d).

Assigning the quantified peptides into domains (rather than proteins) revealed fold-types with strikingly different levels of intrinsic refoldability (Fig. 3i, Extended Data Fig. 5e). Satisfyingly, fold-types exemplified by model proteins are typically efficient refolders (e.g., ribonuclease H-like^28^ (30% non-refoldable), immunoglobulin-like β sandwich domains^29^ (0% non-refoldable), and β-lactamase-like domains (0% non-refoldable). On the other hand, less-characterized folds display markedly higher levels of non-refoldability (e.g., ferritin-like (75% non-refoldable) and class II aaRS-like (80% non-refoldable)).

Proteins of greater molecular weight and essential proteins were also found to be less refoldable (Extended Data Fig. 6a,g). However, these observations are confounded by the fact that, on average, we quantify more peptides for massive proteins and essential proteins, potentially introducing a bias that makes it ‘easier’ to detect a significant structural difference (Extended Data Fig. 6c,i) within them. Importantly, this bias does not affect the other key trends noted (Extended Data Figs. 5b, 6f, 7c, Supplementary Note 2). Alternatively, by lumping together all peptides associated with a given attribute (without regard to which specific protein they came from), this bias vanishes (Supplementary Data), and the key trends we observe are maintained and *more* statistically significant under this bias-free peptide-level analysis (Extended Data Figs. 5a,e, 6e, 7b,e).

Unexpectedly, proteins with cofactors were not generally worse refolders, suggesting that apo-proteins are generally adept at refolding into an intermediate that can select the correct cofactor from a complex mixture (Extended Data Fig. 7d-e). Assigning proteins into cellular compartments and cellular functions shows ribosomal proteins and those involved in translation are the poorest refolders, whilst soluble portions of proteins localized to the inner membrane are surprisingly efficient refolders (Extended Data Fig. 7a-c,f; Supplementary Note 3). Virtually all of these trends were recapitulated in separate experiments in which the proteome was refolded out of urea (Extended Data Fig. 8d-l), and most of the proteins quantified in both the urea and GdmCl experiments had the same refoldability status (Extended Data Fig. 8m-n, Supplementary Data).

### Kinetics of Refolding Across the Proteome

We found that most proteins that *can* refold do so within the first minute (Fig. 4a, Extended Data Fig. 9c). This is consistent with the corpus of classical protein folding studies, which typically record folding transitions on the ms–s timescales^30,31^. Nevertheless, we identified 130 ‘slow’ refolders whose structural dynamics can be probed on the timescale of this experiment (Extended Data Fig. 9e-f). One example is triose phosphate isomerase (TpiA), which is refoldable, however the head of the protein barrel that abuts the active site refolds on the minute-timescale, as observed by the decay kinetics monitored at four independent sites (Fig. 4b). This finding is consistent with previous work that showed TpiA regains its catalytic activity on the minute-timescale^32^. Overall, the refolding proteome was more susceptible to PK after 1 minute of refolding (Extended Data Fig. 9a-b) and as time progressed up to 2 h, it became more native-like (Extended Data Fig. 9c). We could identify 3569 distinct sites which over time became more native-like (|log_2_ (Refolded / Native)| decreased toward zero), of which the majority (64%) became native between 1–5 min (Extended Data Fig. 9d, Supplementary Note 4).

**Fig. 4.**
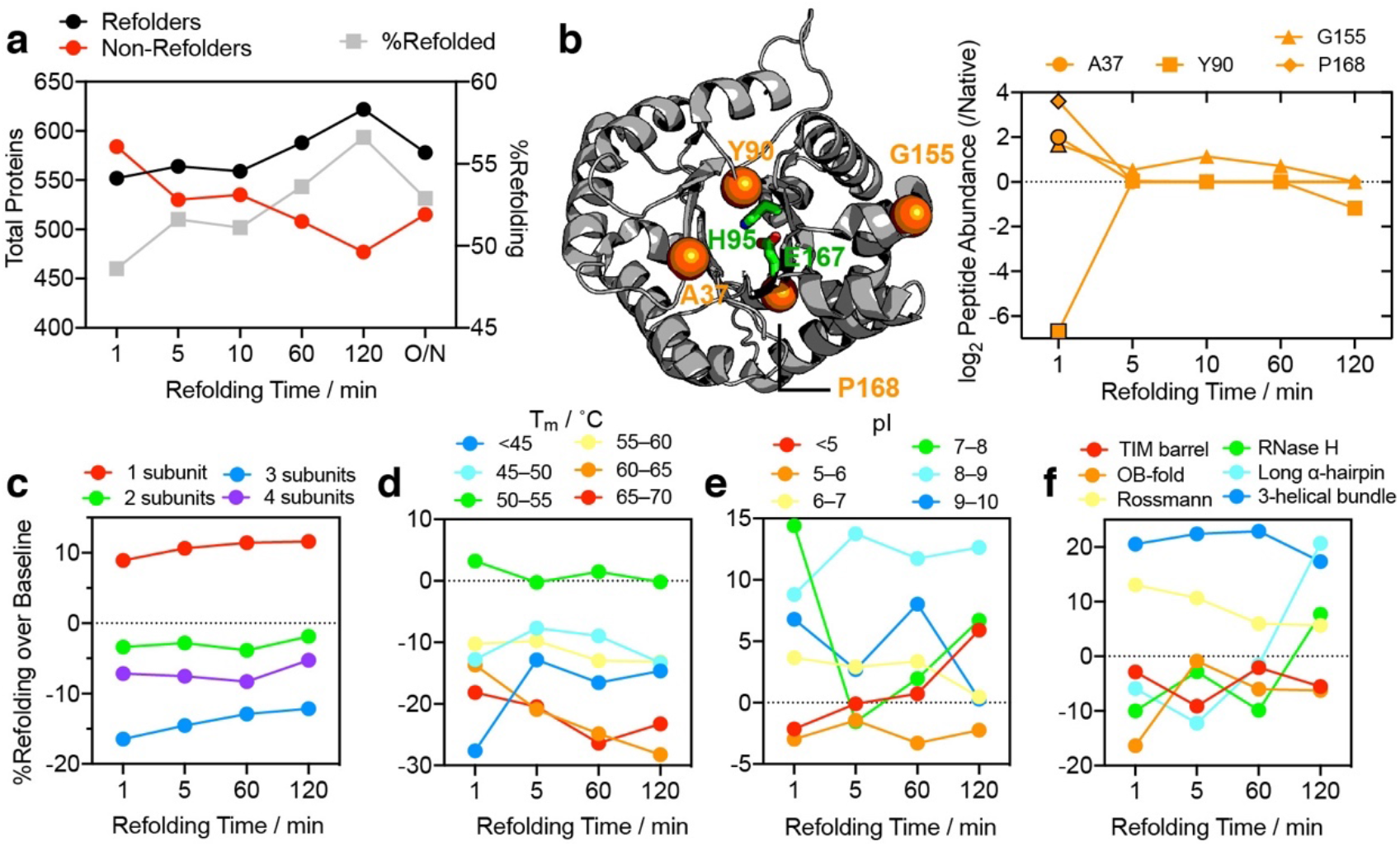
Proteome Refolding Kinetics. **a**, From 1 min to 2 h, the number of refolding proteins increases. Overnight incubation may lead to increased degradation. **b**, Structure of triose phosphate isomerase (TpiA, *at left*) with the positions of four residues (orange) whose accessibility change significantly between 1 min and 2 h (*at right*). All of these residues change conformation in a minute-timescale process associated with forming one side of the TIM-barrel (black rims signify a statistically significant difference relative to native TpiA; P-value < 0.01 by Welch’s t-test). Green residues represent the catalytic residues. **c-f**, Variation in refoldability as a function of time for several classifications: protein subunit count, **c**; midpoint melting temperature in °C, **d**; protein isoelectric point, **e**; and fold-type, **f**. To highlight differences, the percent of refolding proteins (domains) is reported as a difference compared to the overall refoldability of the proteome at each time point. **c**, Dimers and tetramers do not generally require more time to refold, though trimers do. **d**, Proteins with low thermostability tend to refold slowly. **e**, Polyanionic proteins (pI < 5) tend to refold slowly; proteins with pI between 7-8 tend to refold rapidly. **f**, Rossman folds and 3-helical bundles tend to refold rapidly; ribonuclease H-like domains, OB-folds, and long α-hairpins tend to refold slowly. TIM barrels are poorer refolders on all timescales.

Unexpectedly, multimeric proteins were not substantially slower at refolding compared to their monomeric peers at the min-timescale (Fig. 4c, Extended Data Fig. 9h). This suggests that the translational entropy loss associated with refolding multimers is not a global determinant for refolding kinetics on the min-timescale. On the other hand, we found several intriguing relationships between refolding kinetics and thermostability (Fig. 4d), isoelectric point (pI; Fig. 4e), and fold-type (Fig. 4f). Proteins with low pI (<5) are polyanionic and generally refold slowly (Fig. 4e, Extended Data Figs. 9f,j, 10b,d), likely because of the internal charge repulsion within the chain.

Proteins with pI close to cellular pH (7 < pI < 8) have a high propensity to fold very rapidly: this phenotype could have evolved as a defense mechanism against aggregation because charge-neutral proteins are the most prone to aggregate^33^. Proteins with low thermostability (*T*_m_ < 50°C) also had a strong propensity to refold slowly (Fig. 4d, Extended Data Figs. 9f,l, 10f,h,i), which would argue that downhill folding is rare, at least among *E. coli* proteins^34^. We also detected a number of examples (Extended Data Fig. 10d,f,h,i) and an enrichment (Extended Data Fig. 9f) for proteins harbouring divalent cations to refold slowly, suggesting that uptake of these metals could be a rate-limiting step for protein folding on the min-timescale.

Finally, we found that different fold-types showed strikingly distinct kinetic behaviors. Ribonuclease H-like domains, OB-folds, and long alpha-hairpins tend to refold slowly, whereas Rossmann folds and 3-helical bundles have generally completed refolding within one minute (Fig. 4f, Extended Figs. 9m-n, 10d,i). TIM barrels were generally poorer refolders than other fold-types, particularly after given 5 min of refolding time.

### Role of Chaperonins and Thermostability

Chaperones such as the *E. coli* chaperonin GroEL/GroES are known to play indispensable roles in facilitating protein folding *in vivo*^35–37^. However, we found that class III proteins (defined as those with a high propensity to be substrates for GroEL/GroES) were *not* significantly more likely to be non-refoldable (52%) than average (Fig. 5a)^35^. On the other hand, class I proteins (those which are *not* substrates for GroEL/GroES) were almost always non-refoldable (88%; P = 0.06 by chi-square test (5 × 10^−7^ peptide-level)).

**Fig. 5.**
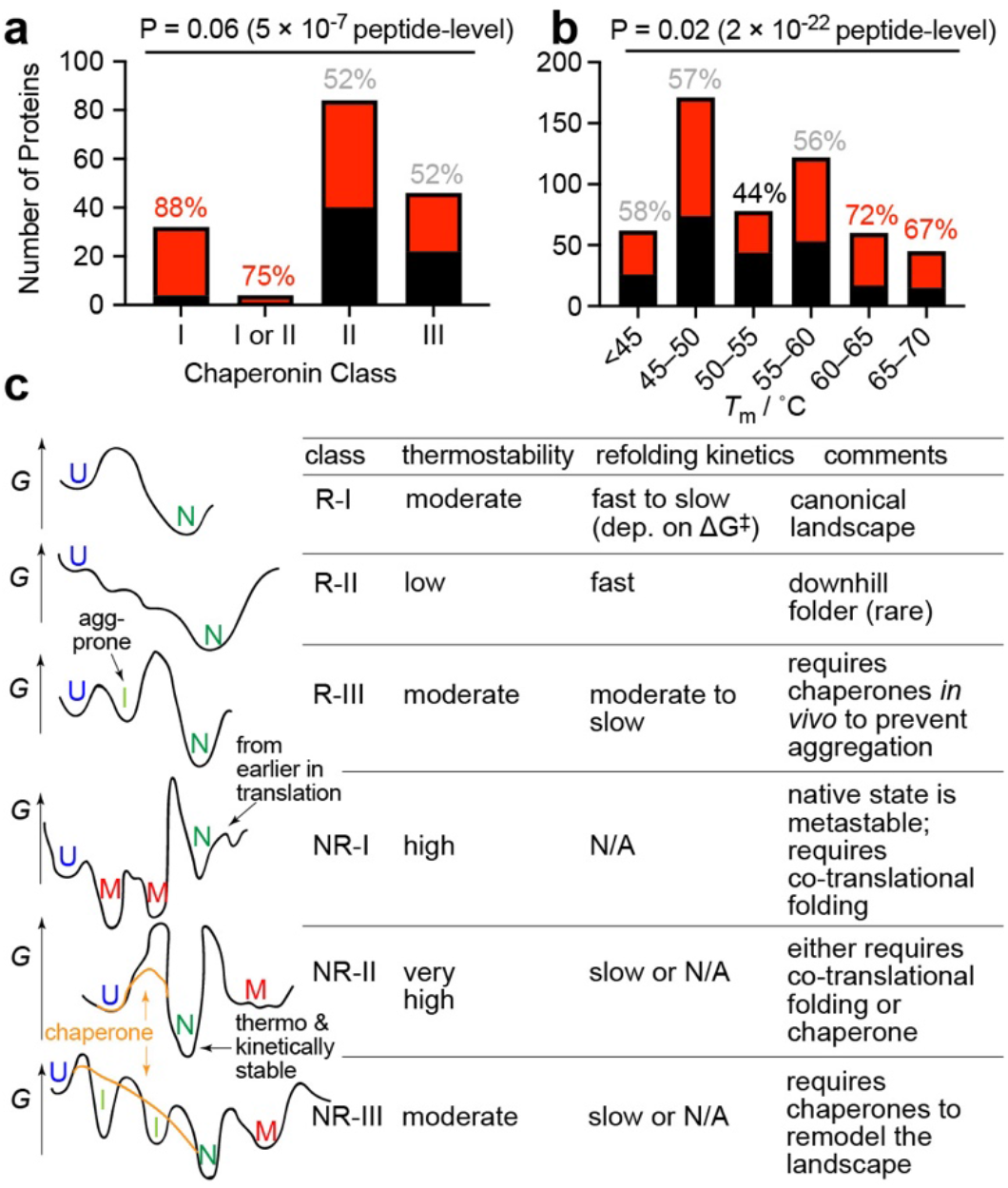
A Taxonomy of Protein Free Energy Landscapes. **a-b**, Number of refoldable (black) and non-refoldable (red) proteins in terms of **a**, chaperonin class^35,36^, and **b**, midpoint melting temperature, *T*_m_^15^. Percentages denote percent of proteins that are non-refoldable (red, higher than average; black, lower than average; gray, average). **a**, Class I proteins are primarily non-refoldable, whilst class II and III proteins are nearly equally distributed. **b**, Highly thermostable proteins are generally non-refoldable, whereas proteins with intermediate thermostability are the most refoldable. Due to low counts, statistical significance is drastically enhanced by conducting analysis at the peptide level. **c**, Six classes of protein free energy landscapes. *G*, Gibb’s free energy; U, unfolded state; N, native state; I, on-pathway intermediate; M, misfolded state; R-, a refoldable class; NR-, a non-refoldable class.

To explain these findings, we speculate that there are two types of class III proteins^35,36^: those whose energy landscapes require tempering by GroEL/GroES to permit access to native states (class NR-III; Fig. 5c), and those which can refold in principle (and do, in our experiments), but do so slowly or via intermediates that are aggregation-prone, hence requiring their sequestration for efficient folding in the cytosol (class R-III). The prior observation that TIM barrels are enriched amongst class III proteins is consistent with our findings that these domains have a higher propensity to either not refold at all or to refold slowly over minutes (Fig. 4f), a timescale over which aggregation might occur in crowded environments.

Comparing our data to estimated midpoint melting temperatures (*T*_m_) for 538 *E. coli* proteins yielded the striking result that the most thermostable proteins (*T*_m_ > 60°C) were preponderantly non-refoldable (70%; P = 0.02 by chi-square test (2 × 10^−22^ at peptide-level)), whereas proteins with intermediate levels of thermostability (50°C < *T*_m_ < 55°C) were the most refoldable (Figs. 5b, 4f). This finding indicates that *E. coli* proteins do not typically achieve thermostability by burrowing their native states into deep free energy wells (which would allow for facile refolding), but instead by ‘protecting’ the native states with steep free energy barriers (class NR-II). These barriers would then prevent re-entry after unfolding. We found further support for this notion by finding that most members (70%) of a cohort of ‘kinetically stable’ proteins that can resist SDS-denaturation are non-refoldable (P = 7 × 10^−4^ by Fisher’s exact test)^38^. Hence, we suggest that a general property of non-refoldable proteins may be that they have very slow unfolding rates, which would imply that cells would not need them to refold after they fold for the first time on the ribosome.

### A New Taxonomy of Protein Free Energy Landscapes

Based on our studies, we propose a taxonomy of protein free energy landscapes that includes three classes of non-refoldable proteins (Fig. 5c). Class NR-II proteins have native states that are thermodynamically stable but surrounded by high barriers, and are defined by high thermostability. Class NR-III proteins must traverse several intermediates separated by sizable barriers to find their native states, and likely require chaperones to smooth their free energy landscapes to fold. Finally, we posit the existence of a third type of non-refoldable protein, class NR-I, which is categorized by a native state that is not thermodynamically stable, but is nevertheless persistent due to steep barriers retaining the full-length chain in a particular region of the landscape.

Class NR-I proteins would rely on co-translational folding (or a pro-domain^12^) to ‘seed’ them in the native region of the energy landscape, where they are then trapped after the protein is fully matured. Such proteins would not be easily rescued by chaperones were they ever to wander away from their native states, and would require degradative mechanisms to maintain proteostasis^37^. We speculate that many class I proteins (non-GroEL/GroES substates) may be of this kind: by folding on the ribosome to a persistent native state, they would not require the service of chaperonins in the cell. *Ipso facto*, they cannot refold after full denaturation of the full-length chain. Multi-subunit assemblies are more likely to be class NR-I because of their high levels of non-refoldability (Fig. 3d), the known role translation plays in their assembly^20–23^, and their dearth among GroEL/GroES substrates^35^. Further refolding experiments in the presence of chaperones and degradation machines will enable the assignment of each non-refoldable protein to these specific sub-classes.

To conclude, we found that many *E. coli* proteins – especially those with many subunits or particular domains – cannot relocate their native states from fully-denatured states solely by following a free energy gradient. We speculate that the free energy landscapes of such full-length proteins may be populated with local (or even global) minima that correspond to non-native soluble structures, which might only be avoidable through processes with kinetic control (e.g., translation). In such a scenario, native states of many proteins may not be thermodynamically stable but kinetically *persistent*^12,38^. By employing a mass spectrometry-based proteomics approach, we have greatly expanded the number of proteins whose refolding has been interrogated, and have shown that many proteins do not follow the same paradigm as most extensively-studied model systems^39^.

## Methods

### Culture and Lysis of K12

*E. coli* cells, strain K12 (NEB ER2738) from saturated overnight cultures, were inoculated in 3 × 50 mL (biological triplicates) of MOPS EZ Rich Defined Media (M2105 Teknova) in 250 mL flasks at a starting OD_600_ of 0.05. Cells were grown at 37 °C with agitation (220 rpm) to a final OD_600_ of 0.8, at which point cells were pelleted by centrifugation at 4000 *g* for 15 mins at 4°C. Supernatants were removed and cell pellets were stored overnight at –20°C until further use.

Cell pellets were resuspended in 1.5 mL of native buffer (20 mM Tris pH 8.0 or 20 mM HEPES pH 7.0, 100 mM NaCl, 2 mM MgCl_2_). Resuspended cells were flash frozen by slow drip over liquid nitrogen and then cryogenically pulverized with a freezer mill (SPEX Sample Prep) over 8 cycles consisting of 1 min grind, 9 Hz and 1 min cool. Pulverized lysates were transferred to a 50 mL centrifuge tube and thawed at room temperature for 20 mins. Cellular lysates were then transferred to a fresh 1.5 mL microfuge tube and clarified at 16000 *g* for 15 mins at 4°C to remove insoluble cell debris. The supernatants were transferred to a fresh microfuge tube and protein concentrations of the clarified cellular lysates were determined by using the bicinchoninic acid assay (Rapid Gold BCA Assay, Pierce) in a microtiter plate format with a plate reader (Molecular Devices). Using the results from the BCA Assay, the clarified cellular lysates were normalized to a protein concentration of 3.3 mg mL^−1^. This was the starting point for most further workflows *vide infra*.

### Precipitation Studies

For the study of protein aggregation, normalized lysates were prepared as described above and carried through various conditions accordingly (see fig. S3). The native sample was prepared by diluting lysate with native buffer (either 20 mM Tris pH 8.0 or 20 mM HEPES pH 7.0, 100 mM NaCl, 2 mM MgCl_2_), either supplemented with 1 mM DTT (or not) to a final protein concentration of 0.23 mg mL^−1^ and incubated overnight at 4°C.

The samples which were refolded by dilution from denaturant were prepared as follows: 600 *μ*L of lysates, solid denaturant (80 mg urea or 100 mg guanidium chloride (GdmCl)), and 2.4 *μ*L of a freshly prepared 700 mM dithiothreitol (DTT) stock solution were added to a fresh 1.5 mL microfuge tube and solvent was removed using a vacufuge plus (Eppendorf) to a final volume of 170 *μ*L, such that the final concentrations of all components were: 11.6 mg mL^−1^ protein, 6M GdmCl or 8 M Urea, 10 mM DTT. Denatured samples were left to unfold at room temperature for 2 h prior to refolding. Unfolded lysates were then diluted 50× with native buffer (either Tris pH 8.0 or HEPES pH 7.0; either supplemented with 1 mM DTT or not) or 500× with native buffer (either Tris pH 8.0 or HEPES pH 7.0; either supplemented with 0.12 M GdnHCl or 0.16 M Urea; either supplemented with 1 mM DTT or not) and incubated overnight at 4°C. After these procedures, the protein concentration will be 0.23 mg mL^−1^ (Rn) or 0.023 mg mL^−1^. This generates 8 refolding conditions carried in biological triplicates (see fig. S3).

The samples which were refolded by dialysis from denaturant were prepared as follows: 35 *μ*L of lysates were diluted with 380 *μ*L of unfolding buffer (20 mM Tris pH 8.0, 100 mM NaCl, 2 mM MgCl_2_, 10 mM DTT) supplemented with either 9 M Urea or 6.6 M GdmCl, such the final concentrations of all components were: 0.28 mg mL^−1^ protein, 6 M GdnHCl or 8 M Urea, 10 mM DTT. Denatured samples were left to unfold at room temperature for 2 h prior to refolding. Unfolded lysates were then transferred to a wetted 3.5K MWCO dialysis cassette according to manufacturer protocol (Slyde-A-Lyzer G2, ThermoFisher) and refolded overnight in native buffer (20 mM Tris pH 8.0, 100 mM NaCl, 2 mM MgCl_2_, supplemented with 1 mM DTT) at 4°C with gentle stirring. After anticipated swelling of the dialysate, the protein concentration is expected to be 0.23 mg mL^−1^. This generates 2 refolding conditions carried in biological triplicates (see fig. S3).

The samples which were refolded by slow cooling from thermal denaturation were prepared as follows: 14 *μ*L of lysates were diluted with native buffer (either Tris pH 8.0 or HEPES pH 7.0, 100 mM NaCl, 2 mM MgCl_2_, supplemented with 1 mM DTT) to a final protein concentration of 0.23 mg mL^−1^ or 0.023 mg mL^−1^ and distributed 100 *μ*L per PCR tube. Samples were heated to 90°C over 1 hr and then slowly cooled overnight to 4 °C (−1°C per 20 minutes) using a Thermocycler (ProFlex PCR System, ThermoFisher). Refolded samples were resuspended through pipetting and combined into a fresh 1.5 mL microfuge tube. This generates 3 refolding conditions carried in biological triplicates (see fig. S3).

Native and refolded lysates were clarified at 16000 *g* for 1 hr at 4°C. The supernatant was removed by careful pipetting so as not to disturb the protein pellet. The pellet was then resuspended in resuspension buffer (100 *μ*L 50 mM Ammonium Bicarbonate, 8 M urea) and the protein concentration was determined by BCA Assay as described above. The protein concentrations were converted to amounts using the constant volume and then converted to a fractional precipitation using the constant initial loading (46 *μ*g of protein). The data are reported as a means ± standard deviations from the biological triplicates which were differentiated at the inoculation stage. Statistical tests were carried out using ANOVA with follow-up pairwise tests using Tukey’s correction for multiple hypothesis testing, as implemented in Prism 8 (Graphpad). The “precipitation” measured for the native samples were treated as background of the experiment because there should not be any precipitated protein in them. This measurement was done either with or without 1 mM DTT to correct for the known background signal it generates in the BCA assay.

Dynamic Light Scattering (DLS) studies were performed as follows: 100 *μ*L of each of the samples refolded at 0.23 mg mL^−1^, 200 *μ*L of each of the samples refolded at 0.023 mg mL^−1^, and 100 *μ*L of a 5× dilution in native buffer of each of the thermal refolded samples were transferred to a black walled 96 well plate (Falcon). Protein sizes were measured using a Dyna Pro Plate Reader II (Wyatt Technologies). Each time-dependent autocorrelation function was acquired over the course of 5 secs, repeated 15 times for each sample well and averaged together at an operating temperature of 25°C. Size distributions of proteins were obtained using the in-built DYNAMICS (Wyatt Technologies) software. Intensities were rebinned according to the size distributions shown in fig. S5, and averaged over biological triplicates.

### Single Protein Experiments (SNase and RNase H)

For the study of single protein refolding, wild-type staphylococcal nuclease (SNase) and wild-type ribonuclease H from *Thermus thermophilus* (*Tt*RNase H) were expressed and purified as described previously^40,41^ and were provided as generous gifts from the García-Moreno Lab at Johns Hopkins University as 10 mg mL^−1^ frozen stocks in water. The native samples were prepared by diluting the frozen protein stock (SNase or *Tt*RNase H) 450× with a 49:1 mixture of native buffer (20 mM Tris pH 8.0, 100 mM NaCl, 2 mM MgCl_2_, 1 mM DTT) and unfolding buffer (20 mM Tris pH 8.0, 100 mM NaCl, 2 mM MgCl_2_, 9 M Urea, 10 mM DTT). The native samples were then concentrated to a final protein concentration of 0.23 mg mL^−1^ using 3K MWC Amicon® Ultra-15 Centrifugal Filter Units (Millipore Sigma) (Amicon Filter). For each protein, we generated a native sample in technical triplicates (see fig. S1 for workflow).

The pure protein samples were unfolded by diluting frozen protein stocks (SNase or *Tt*RNaseH) 9× with unfolding buffer (final protein concentration was 1.1 mg mL^−1^ and final concentration of urea was 8 M) and incubating for 1 h at room temperature. Unfolded proteins were then refolded by either dilution or dialysis. To prepare the samples that were refolded by dilution, unfolded proteins were diluted 50× with native buffer and incubated at room temperature for 120 m to allow for the protein to refold. The samples was then concentrated to a final protein concentration of 0.23 mg mL^−1^ using Amicon Filters according to manufacturer protocol. These two samples were further analyzed by circular dichroism. To prepare the samples that were refolded by dialysis, 115 *μ*L unfolded proteins were diluted with 300 *μ*L of unfolding buffer such that the final concentration of all components were 0.3 mg mL^−1^ protein, 8 M Urea, 10 mM DTT. Unfolded proteins were then transferred to a wetted 3.5K MWCO dialysis cassette according to manufacturer protocol (Slyde-A-Lyzer G2, ThermoFisher) and refolded overnight in native buffer at 4°C with gentle stirring. After anticipated swelling of the dialysate, the protein concentration is expected to be 0.23 mg mL^−1^.

Circular Dichroism (CD) studies of protein folding were conducted as follows: protein concentration of each refolded sample was determined by measuring the absorbance at 280 nm and calculating the protein concentration using the extinction coefficients of the proteins (15,930 M^−1^ cm^−1^ for SNase and 30,480 M^−1^ cm^−1^ for *Tt*RNaseH). CD spectra of refolded proteins were obtained at 25°C using a CD Spectrometer (Aviv 420) over a spectral range of 198 nm to 250 nm at a scanning rate of 1 nm / 3 sec in a 1-mm pathlength cuvette. Molar Residue Ellipticity (MRE) was calculated for the minima of each of the refolded proteins (222 nm for SNase and 215 nm for *Tt*RNaseH) using the protein concentration and the number of amino acids in each protein.

200 *μ*L each of the refolded (dilution or dialysis) samples were divided into two different aliquots, in which one was spiked with 14.5 *μ*L concentrated clarified *E.coli* lysate (~3.3 mg mL^−1^) to a final concentration of 0.23 mg mL^−1^ or not. The samples were then incubated with proteinase K (PK) from *Tritirachium album* (Sigma Aldrich).

2 *μ*L of a PK stock (prepared as a 0.25 mg mL^−1^ PK in a 1:1 mixture of native buffer and 20% glycerol, stored at −20°C and only thawed once) were added to a fresh 1.5 mL microfuge tube. 200 *μ*L of the native or refolded proteins (pure or spiked with lysate) were then added to the same microfuge tube and rapidly mixed by pipetting (enzyme: substrate ratio of 1:100 on a weight basis). Samples were incubated for exactly 1 min in a water bath preequilibrated at 25°C before transferring them to a mineral oil bath preequilibrated at 100°C and incubating them for 5 mins to quench PK activity. Boiled samples were then transferred to a fresh 2 mL microfuge tube containing 200 mg urea and 85 *μ*L of native buffer such that the final urea concentration was 8 M and the final volume was 415 *μ*L.

All protein samples were prepared for mass spectrometry as follows: 6 *μ*L of a freshly prepared 700 mM stock of DTT were added to the sample containing microfuge tube to a final concentration of 10 mM and incubated at 37°C for 30 minutes at 700 rpm on a thermomixer to reduce cysteine residues. 24 *μ*L of a freshly prepared 700 mM stock of iodoacetamide (IAA) were then added to a final concentration of 40 mM and incubated at room temperature in the dark for 45 minutes to alkylate reduced cysteine residues. After alkylation of cysteines, 1215 *μ*L of 100 mM ammonium bicarbonate were added to the samples to dilute the urea to a final concentration of 2 M. 1 *μ*L of a 1 mg mL^−1^ stock of Trypsin (Pierce) was added to the samples (to a final enzyme:substrate ratio of 1:50 on a weight basis) and incubated overnight at 25°C at 700 rpm.

Peptides were desalted with Sep-Pak C18 1 cc Vac Cartridges (Waters) over a vacuum manifold. Tryptic digests were first acidified by addition of 16.6 *μ*L trifluoroacetic acid (TFA, Acros) to a final concentration of 1% (vol/vol). Cartridges were first conditioned (1 mL 80% ACN, 0.5% TFA) and equilibrated (4 × 1 mL 0.5% TFA), before loading the sample slowly under a diminished vacuum (ca. 1 mL/min). The columns were then washed (4 × 1 mL 0.5% TFA), and peptides eluted by addition of 1 mL elution buffer (80% ACN, 0.5% TFA). During elution, vacuum cartridges were suspended above 15 mL conical tubes, placed in a swing-bucket rotor (Eppendorf 5910R), and spun for 3 min at 350 *g*. Eluted peptides were transferred from Falcon tubes back into microfuge tubes and dried using a vacuum centrifuge (Eppendorf Vacufuge). Dried peptides were stored at −80°C until analysis. For analysis, samples were vigorously resuspended in 0.1% FA / 2% ACN in Optima water (ThermoFisher) to a final concentration of 1 mg/mL.

### Kinetic Refolding Experiments from GdmCl and Urea

For the study of proteome refolding kinetics, normalized lysates were prepared as described above. The native samples (N) were prepared by diluting lysates with native buffer (20 mM Tris pH 8.5, 100 mM NaCl, 2 mM MgCl_2_) supplemented with 1 mM DTT to a final protein concentration of 0.23 mg mL^−1^ and incubated overnight at 4°C.

The samples which were refolded by dilution from denaturant (R) were prepared as follows: 600 *μ*L of lysates, solid denaturant (80 mg urea or 100 mg guanidium chloride (GdmHCl)), and 2.4 *μ*L of a freshly prepared 700 mM dithiothreitol (DTT) stock solution were added to a fresh 1.5 mL microfuge tube and solvent was removed using a Vacufuge plus (Eppendorf) until a final volume of 170 *μ*L, such that the final concentrations of all components were: 11.6 mg mL^−1^ protein, 6M GdmCl or 8 M urea, 10 mM DTT. Denatured samples were left to unfold at room temperature for 2 h prior to refolding. To refold unfolded lysates, 4 *μ*L of unfolded lysates were transferred to a fresh 1.5 mL microfuge tube and then diluted 50× with native buffer supplemented with 1 mM DTT and such that the final protein conc. was 0.23 mg mL^−1^. Refolded lysates were incubated at room temperature for different durations (1 m, 2 m, 5 m, 10 m, 60 m, 120 m, or overnight) to allow for proteins to refold before incubation with PK (see Fig. 1).

2 *μ*L of a PK stock (prepared as a 0.25 mg mL^−1^ PK in a 1:1 mixture of native buffer and 20% glycerol, stored at −20°C and only thawed once) were added to a fresh 1.5 mL microfuge tube. After allowing proteins to refold for a specified amount of time, 200 *μ*L of the refolded lysates were then added to the same microfuge tube and rapidly mixed by pipetting (enzyme: substrate ratio is 1:100 on a weight basis). Samples were incubated for exactly 1 min in a water bath preequilibrated at 25°C before transferring them to a mineral oil bath preequilibrated at 100°C and incubating them for 5 mins to quench PK activity. Boiled samples were then transferred to a fresh 2 mL microfuge tube containing 200 mg urea and 85 *μ*L of native buffer such that the final urea concentration was 8 M and the final volume was 415 *μ*L. This method generates the limited proteolysis sample (LiP; further abbreviated as L) protein samples. For samples designated as controls (C), the same procedure was used as above, except PK was not added. For samples designed as native (N; native control (NC) as well as native LiP (NL)), samples were prepared as above, except they were not unfolded and refolded. In total, 30 samples were prepared for this experiment; native control, native LiP, refolded Control, each done in biological triplicates. The refolded LiP were generated for each of the seven refolding times in biological triplicates.

All protein samples were prepared for mass spectrometry and desalted with Sep-Pak C18 1cc Vac Cartridges (Waters) exactly as described above.

### LC-MS/MS

Chromatographic separation of digests were carried out on a Thermo UltiMate3000 UHPLC system with an Acclaim Pepmap RSLC, C18, 75 *μ*m × 25 cm, 2 *μ*m, 100 Å column. Approximately 1.5 *μ*g of protein was injected onto the column. The column temperature was maintained at 40°C, and the flow rate was set to 0.300 *μ*L min^−1^ for the duration of the run. Solvent A (0.1% FA) and Solvent B (0.1% FA in ACN) were used as the chromatography solvents.

The samples were run through the UHPLC System as follows: peptides were allowed to accumulate onto the trap column (Acclaim PepMap 100, C18, 75 *μ*m × 2 cm, 3*μ*m, 100 Å column) for 10 min (during which the column was held at 2% Solvent B). The peptides were resolved by switching the trap column to be in-line with the separating column, quickly increasing the gradient to 5% B over 5 min and then applying a 95 min linear gradient from 5% B to 25% B. Subsequently, the gradient was increased from 35% B to 40% B over 25 minutes, and then increased again from 40% B to 90% B over 5 minutes. The column was then cleaned with a saw-tooth gradient to purge residual peptides between runs in a sequence.

A Thermo Q-Exactive HF-X Orbitrap mass spectrometer was used to analyze protein digests. A full MS scan in positive ion mode was followed by twenty data-dependent MS scans. The full MS scan was collected using a resolution of 120,000 (@ m/z 200), an AGC target of 3E6, a maximum injection time of 64 ms, and a scan range from 350 to 1500 m/z. The data-dependent scans were collected with a resolution of 15,000 (@ m/z 200), an AGC target of 1E5, a minimum AGC target of 8E3, a maximum injection time of 55 ms, and an isolation window of 1.4 m/z units. To dissociate precursors prior to their re-analysis by MS2, peptides were subjected to an HCD of 28% normalized collision energies. Fragments with charges of 1, 2, and >8 were excluded from analysis, and a dynamic exclusion window of 30.0 s was used for the data-dependent scans.

### MS Data Analysis

Proteome Discoverer Software Suite (v2.4, Thermo Fisher) and the Minora Algorithm were used to analyze mass spectra and perform Label Free Quantification (LFQ) of detected peptides. Default settings for all analysis nodes were used except where specified. The data were searched against *Escherichia coli* (UP000000625, Uniprot) reference proteome database. For peptide identification, the Proteome Discoverer Sequest HT node was using a semi-tryptic search. A precursor mass tolerance of 10 ppm was used for the MS1 level and a fragment ion tolerance was set to .02 Da at the MS2 level. Oxidation of methionine and Acetylation of the N-terminus were allowed as dynamic modifications while carbamidomethylation on cysteines was set as a static modification. Raw extracted ion intensity data for the identified peptides were exported from the .pdResult file using a three-level hierarchy (protein > peptide group > consensus feature). Moreover, a second file listing all consensus features (regardless of whether they were assigned to a peptide group) was exported from the .pdResult file. These raw data were processed utilizing custom Python scripts (see Supplementary Text 1).

To analyze trends in refoldability between different classifications, we compiled together the number of significant peptides and the total number of peptides quantified for each protein (domain) along with various metadata assembled from EcoCyc, the SUPERFAMILY database, and the protein isoelectric database (see Supplementary Text 2). Tests of categorical significance was accomplished primarily using the chi-square test (as implemented in Excel), or in a few cases, Fisher’s exact test when only two groups were being compared. Non-parametric analyses of distributions were conducted in GraphPad Prism 8.

### Bioinformatics and Metadata Collection

Many of our findings depend on assigning a range of metadata to each of the proteins in the *E. coli* proteome and identifying patterns in refoldability between proteins and those classifications. The vast majority of the metadata that we used for these purposes came from the EcoCyc database (http://ecocyc.org), a curated database of the genes, proteins, and metabolic networks in the K-12 strain of *E. coli*^42^.

We used the gene symbol as our main identifier for *E. coli* proteins (abcX) although include a list of all synonyms identified by Ecocyc for that gene to facilitate a cross-comparison to the Uniprot and SUPERFAMILY databases. Ecocyc provides information about cellular compartment (cytosol, inner membrane, periplasmic space, outer membrane, ribosome, cell projection), subunit composition, essentiality, copy number, cofactors, and molecular weight (from nucleotide sequence). When the information was available, we used Ecocyc’s *Component Of* category in order to obtain the full constitutive composition of the protomer within a complex. Cellular compartment, subunit composition, and cofactor information is derived in a manually-curated manner from review of the relevant literature for each protein.

We note that subunit composition information is complicated to define precisely, as many proteins form non-constitutive complexes but those interactions are not required for the protein to be stably folded. We further note that Ecocyc’s collection of cofactor information is imperfect (comparison to PDB structures at time revealed disparities; e.g., LigA has both Mg^2+^ and Zn^2+^ cofactors although Ecocyc annotation includes only Mg^2+^), as well as the imperfect nature of defining a structural cofactors – some cofactors are intrinsic to a protein’s structure whereas others turn over in the manner of a substrate. We have opted to use Ecocyc’s information on record ‘as is’ for reproducibility and consistency, and developed a program that collects this information from the database and inserts it into a file, which is available upon reasonable request.

Copy number information predominantly comes from a single ribosome profiling study by Li and coworkers^43^. We used copy number in Neidhardt EZ rich defined medium because of its similarity to the growth medium used in these studies. Essentiality information predominantly comes from the Keio collection study by Baba and coworkers^44^. We used essentiality in LB (Lennox) media because of its use in the creation of the Keio collection. Abundance on a weight basis was determined by multiplying copy number by molecular weight.

Domain information was based on the SCOP hierarchy, and obtained through the SUPERFAMILY database (http://supfam.org)^45,46^. We used custom scripts to edit the ‘raw’ file available from supfam.org into a format more usable for our purposes (including the switch from a Uniprot identifier to the gene symbol identifier). This file is which is available upon reasonable request.

Isoelectric effects were obtained from the isoelectric database^18^. We downloaded the file corresponding to *E. coli* and took the average of the isoelectric point calculated by all algorithms available for each protein. Midpoint melting temperatures were obtained from Leuenberger et al.^15^. Specifically, we downloaded aai7825_Leuenberger_Table-S3 and used the column entitled “Tm Protein,” a melting temperature based on a hierarchical fitting procedure. Chaperonin classes were obtained from Kerner et al.^35^. Specifically, we examined Table S3 and manually identified the current Uniprot accession code for each of the proteins identified by Kerner et al., and transferred this information into a file that contains the gene symbol, the current Uniprot accession code, and the class assignment. We have also included information from Fujiwara et al.^36^, which breaks down Class III proteins into Class III^−^ and Class IV in our Supplementary Data.

## Data Availability

Raw proteomic data has been submitted to PRIDE via the ProteomeXchange under accession codes XXXXXXX. Data used to construct figures are provided as spreadsheets under Supplementary Data. Python programs are available on GitHub at XXXXX.

## Acknowledgments

We thank S. Marqusee and D. Barrick for thoughtful conversations, and P. Mortimer for support with and maintenance of mass spectrometer instrumentation. We are grateful to B. García-Moreno and co-workers for providing samples of SNase and *Tt*RNase H, to S. Myong and co-workers for assistance with DLS measurements, and to K. Tripp for assistance with CD measurements. We acknowledge HFSP (RGY0074/2019) for funding. B.W. thanks the Program in Molecular Biophysics training grant (NIH T32GM008403) and H.E.T. thanks the Chemistry-Biology Interface training grant for support (NIH T32GM080189).

## Author contributions

S.D.F. designed the experiments with input from P.T. P.T. performed proteomics experiments with assistance from B.W. and H.E.T. P.T. and S.D.F. performed analyses. S.D.F wrote the manuscript with input from all authors.

## Additional information

**Supplementary information** is available at XXXXXX.

**Correspondence and requests for materials** should be addressed to S.D.F.

---

**Extended Data Fig. 1.**
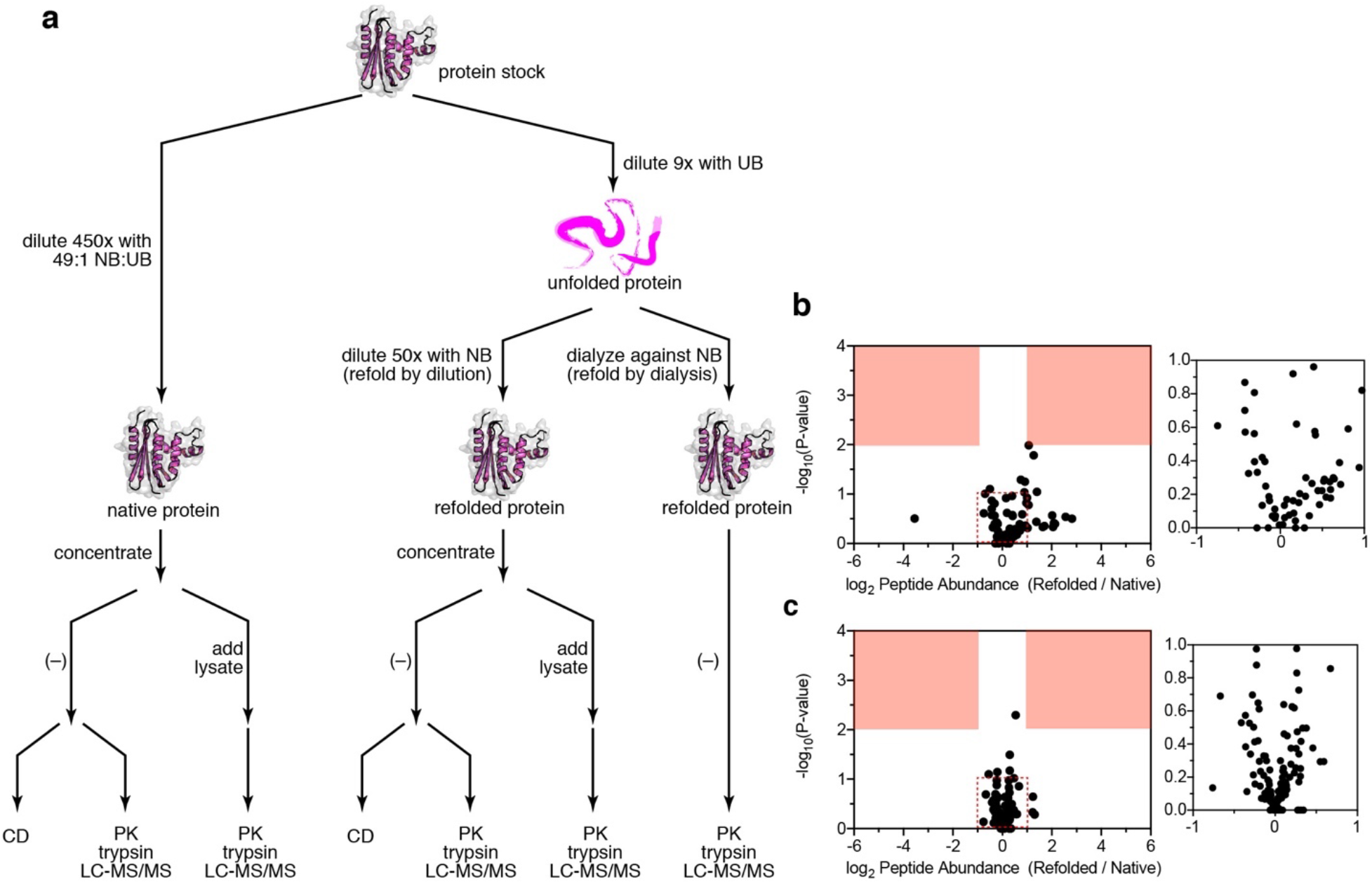
Refolding of Model Proteins. **a**, Workflow for experiments on purified proteins. A concentrated protein stock was either diluted 450-fold with a 49:1 mixture of native buffer (NB) and unfolding buffer (UB) to generate a native sample, or diluted 9-fold with unfolding buffer, incubated, and diluted 50-fold further to generate a refolded sample; see Methods. **b-c**, Volcano plot comparing native proteins to their refolded forms after unfolding and refolded by dialysis. Peptide abundances from native and refolded Staphylococcal nuclease (SNase, **b**), and ribonuclease H from *Thermus thermophilus* (*Tt*RNase H, **c**) (*n* = 3). Effect sizes reported as ratio of averages, and P-values are based on Welch’s test. Red regions designate significance (effect-size > 2, P-value < 0.01). Inset shows large number of points clustered near the origin. The data suggest no significant difference in the structure of native SNase and *Tt*RNase H and the conformation produced when it is refolded by dialysis out of urea.

**Extended Data Fig. 2.**
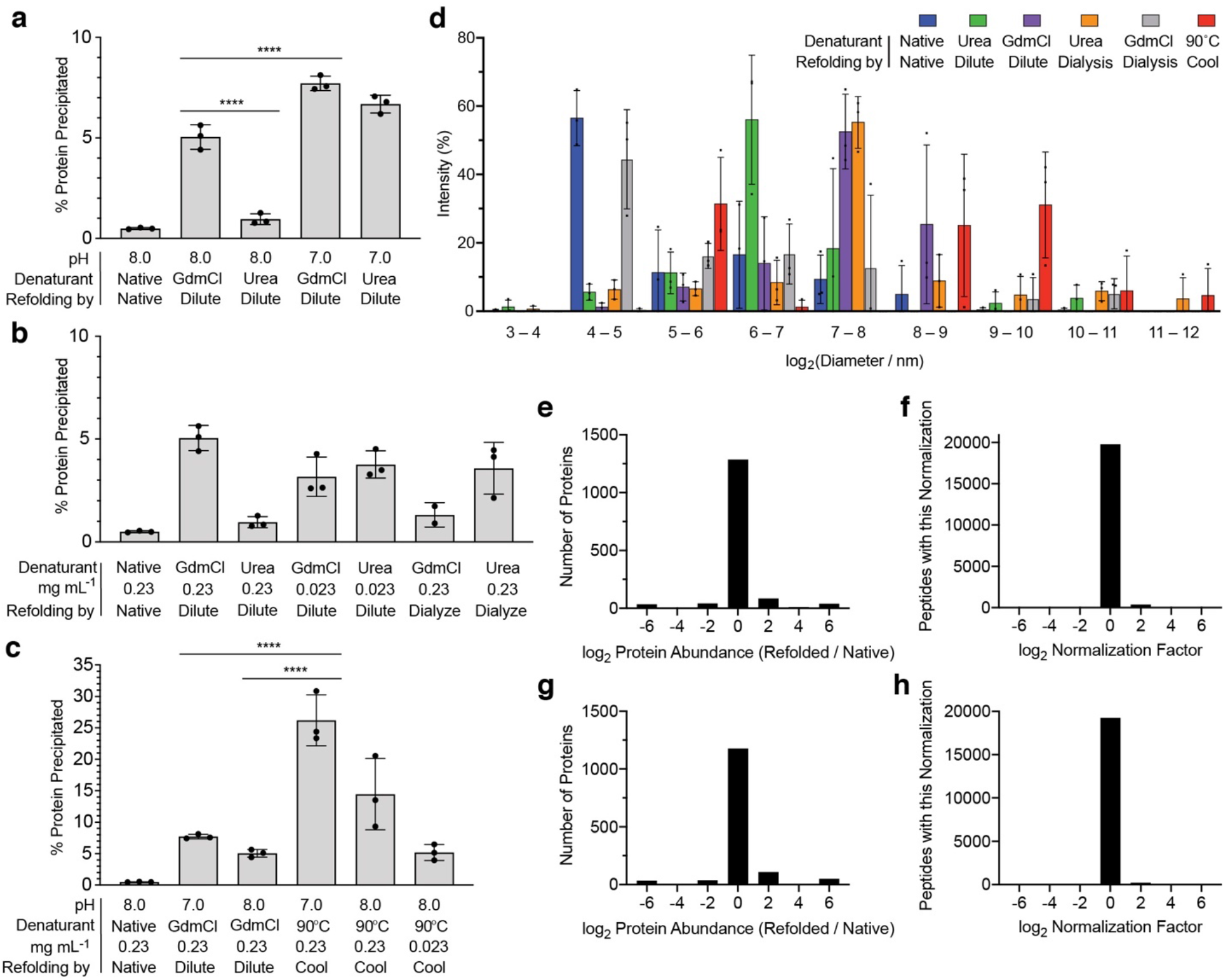
Pelleting Assays and Dynamic Light Scattering Assays to Monitor Aggregation Formed upon Refolding *E. coli* Lysates. **a-c**, Assays to determine conditions which minimize the amount of precipitation formed upon refolding clarified *E. coli* extracts. Precipitation was quantified via the BCA assay (*n* = 3); see Methods. **a**, Comparison of pH and choice of denaturant on protein precipitation levels. Conducting unfolding/refolding at pH 7.0 results in significantly more precipitation than at pH 8.0 (P-value < 0.0001 by ANOVA test with follow-up pairwise comparison with Tukey’s correction). Refolding from GdmCl results in significantly more precipitation than refolding from urea (P-value < 0.0001 by ANOVA test with follow-up pairwise comparison with Tukey’s correction). **b**, Comparison of protein concentration (0.23 mg/mL vs. 0.023 mg/mL) and denaturant removal method (e.g., 50x dilution vs. dialysis overnight) on protein precipitation levels. No significant effects were found, based on ANOVA. **c**, Comparison of various methods to unfold and refold proteins by thermal denaturation followed by slow cooling compared to chemical denaturants. Thermal annealing resulted in significantly more precipitation than chemical denaturants (P-value < 0.0001 by ANOVA test with follow-up pairwise comparisons with Tukey’s correction). This effect could be mitigated by performing thermal unfolding/refolding at pH 8.0 and very low concentration. **d**, Extracts were unfolded and refolded according to the conditions given, and then an aliquot of the mixture was pipetted into a flat-bottom opaque-walled microtiter plate for analysis by DLS (*n* = 3) to quantify soluble aggregates; see Methods. In native extracts (that were never subject to unfolding), particles had an overall distribution with smaller diameters (average diameter = 16.9 ± 4.4 nm (mean ± std. dev. across replicates)). In thermally annealed extracts, particles had an overall distribution with large diameters (average diameter = 783 ± 882 nm (mean ± std. dev. across replicates); 275 ± 66 nm if an outlier is discarded). In extracts subject to chemical denaturation followed by refolding via dilution or dialysis, particle diameter distributions were intermediate between these two extrema. Unfolding by GdmCl followed by refolding via dialysis resulted in particles whose distribution of diameters was similar to native (average diameter = 27.1 ± 10.7 nm (mean ± std. dev. across replicates)). Refolding by dilution generally furnished particles whose distributions of diameters were somewhat larger. **e-h**, Few proteins precipitate under the optimized refolding conditions according to LC-MS/MS. **e-f**, Control samples in which native and GdmCl-refolded *E. coli* lysates were not subjected to limited proteolysis were generated in order to determine overall protein abundance differences between these two samples. **e**, The vast majority of the proteins were present in equal abundance between native and refolded samples. **f**, Each peptide derived from a protein that was present with significantly different abundance between the native and refolded samples (abundance ratio > 2, P-value < 0.01) was adjusted with a normalization factor. A histogram of all 20274 normalization factors shows the vast majority (~99%) were unity. **g-h**, As in **e-f**, except for control samples comparing native and urea-refolded *E. coli* lysate.

**Extended Data Fig. 3.**
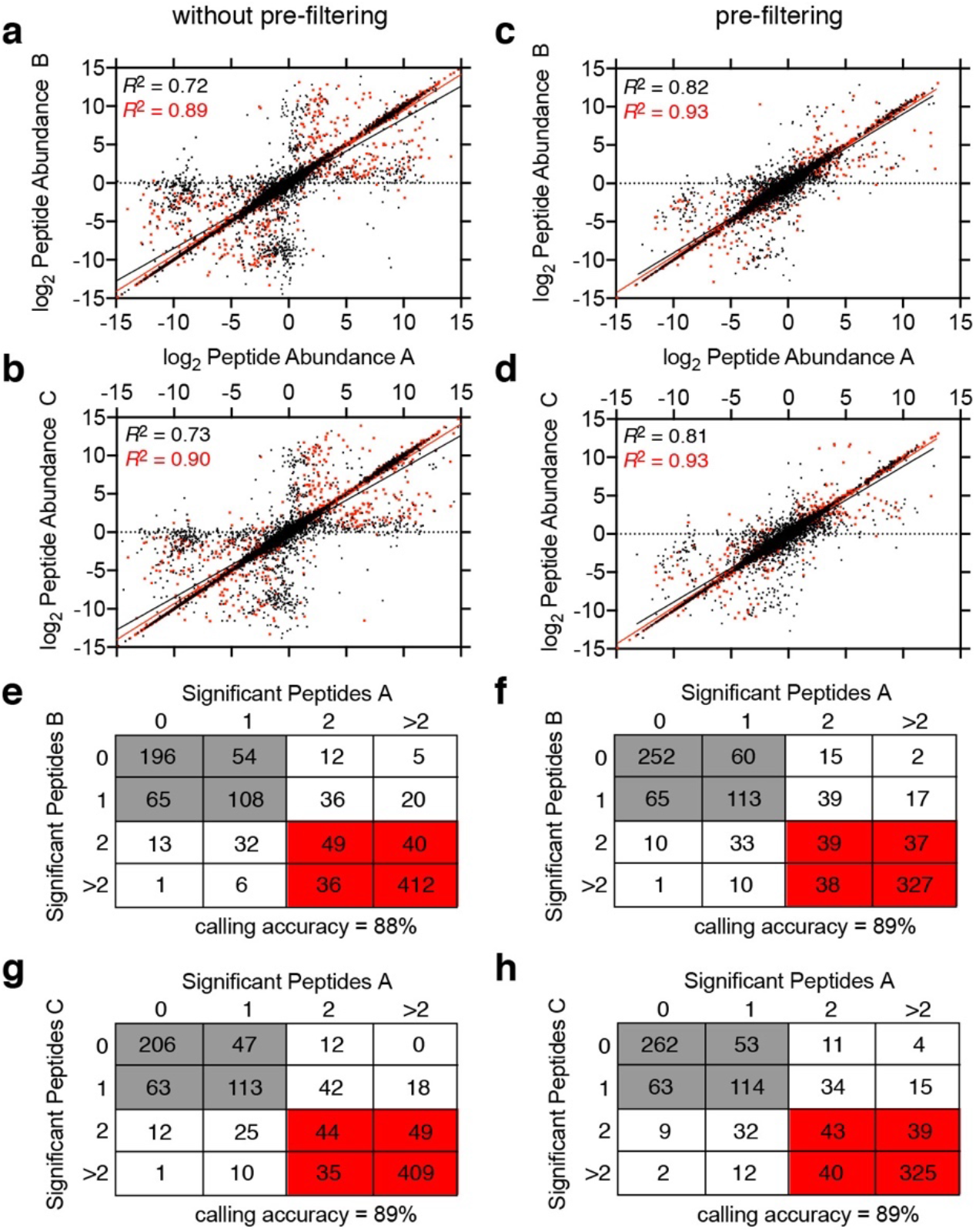
Reproducibility Analysis of Proteome-wide Refolding from GdmCl. **a-b**, The reproducibility in peptide abundance ratios (refolded/native) between three technical replicates of the experiment in which *E. coli* lysate was refolded out of GdmCl overnight. Each of these experiments in turn consisted of three biological replicates of native and three biological replicates of refolded. Black points designate those where the abundance difference between native and refolded was not significant and red points designate those that were significant (P-value < 0.01 by Welch’s t-test). Pearson correlation coefficients are provided; **a**, reproducibility between replicate A and replicate B; **b**, reproducibility between replicate A and replicate C. **c-d**, As in **a** and **b**, except applying a filtering algorithm is applied to test whether peptides with missing data should be retained or dispensed (see Supplementary Text 1). The algorithm successfully removes the majority of extreme values that are not reproducible between technical replicates and significantly improves Pearson correlation coefficients. **e-h**, Calling accuracy matrices showing the number of proteins for which 0, 1, 2, or >2 significant peptides were mapped to a given protein in either replicate for the subset of proteins that were quantified in both replicates A and B, **e-f**; or for the subset of proteins that were quantified in both replicates A and C, **g-h**. Gray boxes correspond to proteins that were called refoldable in both experiments, and red boxes correspond to proteins that were called non-refoldable in both experiments. White boxes correspond to proteins that were called differently in the different replicates. Calling accuracies calculated both without (**e,g**) and with (**f,h**) the filtering algorithm.

**Extended Data Fig. 4.**
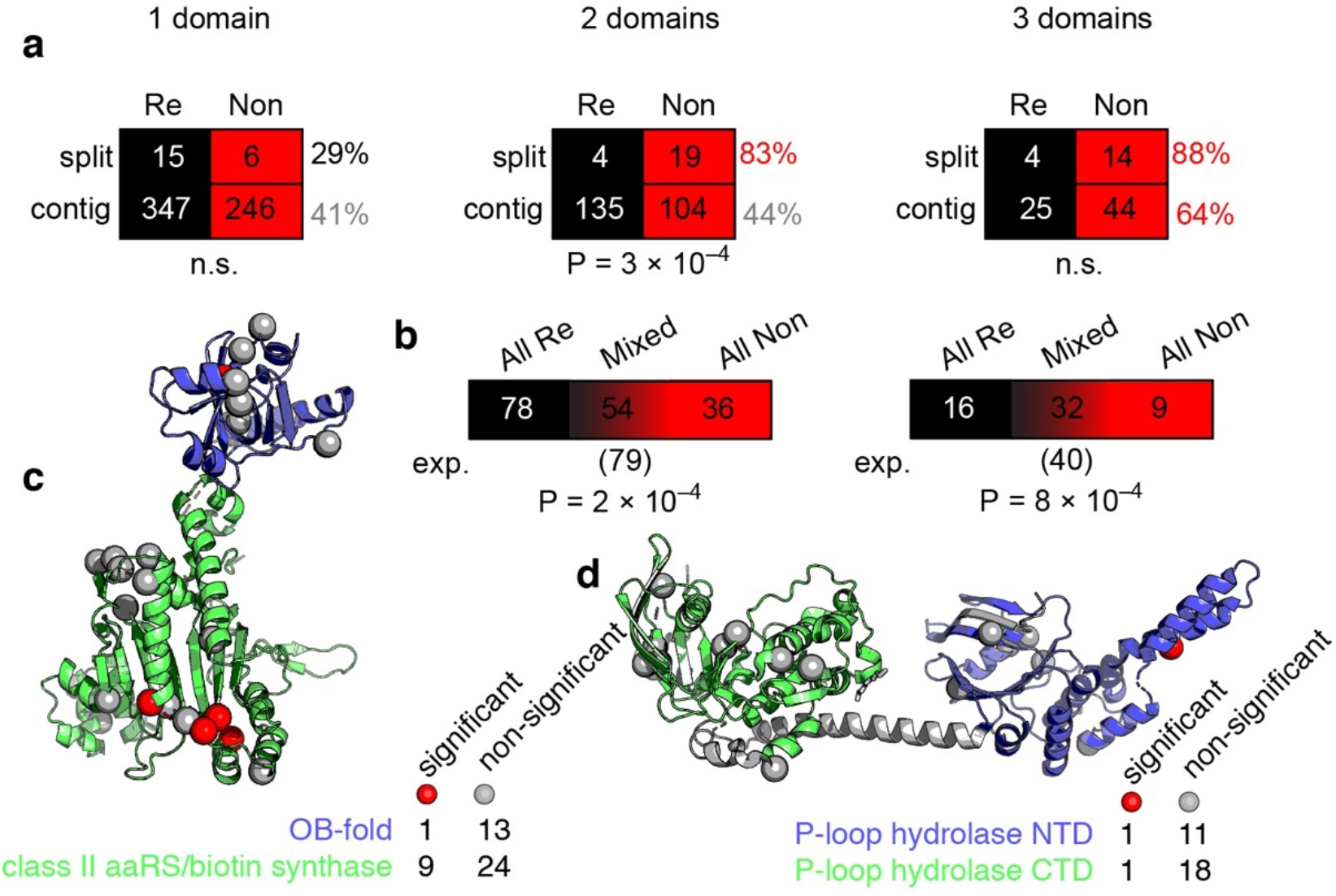
Domain Refolding Coupling. **a**, Splits in domain structure affect refoldability. Contingency tables showing the number of proteins with 1, 2, or 3 domain(s) that are refoldable (or non-refoldable) divided into the classification if the protein has domains that are split into multiple ranges with intervening sequences (split) or contiguous (contiguous), as based on domain ranges from SUPERFAMILY 2. Split 1-domain proteins tend to be *more* refoldable, though the effect is not statistically significant. Split 2-domains are significantly less refoldable (P = 3 × 10^−4^ by Fisher’s exact test) than 2-domain proteins in which the domains are contiguous. Split 3-domain proteins are somewhat less refoldable than 3-domain proteins in which the domains are contiguous (though all 3-domain proteins are generally non-refoldable). **b**, Contingency tables showing the number of 2- or 3-domain proteins in which all domains are refoldable, all domains are non-refoldable, or a mixture. The refolding fates of separate domains on the same protein tend to be coupled; the number of proteins in which the domains have different refolding fates (i.e., mixed) is smaller than what would be expected based on the binomial distribution. P-values based on chi-square test. **c**, X-ray structure of lysyl-tRNA synthetase (LysS; PDB: 1BBW), a 2-domain protein in which one domain (an OB-fold) is refoldable and the other (a class II aaRS/biotin synthase-like domain) is not. Note that the latter domain is almost always non-refoldable (see Fig. 3i). **d**, X-ray structure of energy-dependent translational throttle protein (EttA; PDB: 4FIN), a 2-domain protein in which both domains are refoldable. This scenario is significantly more common. Only half-tryptic sites are mapped to the structures.

**Extended Data Fig. 5.**
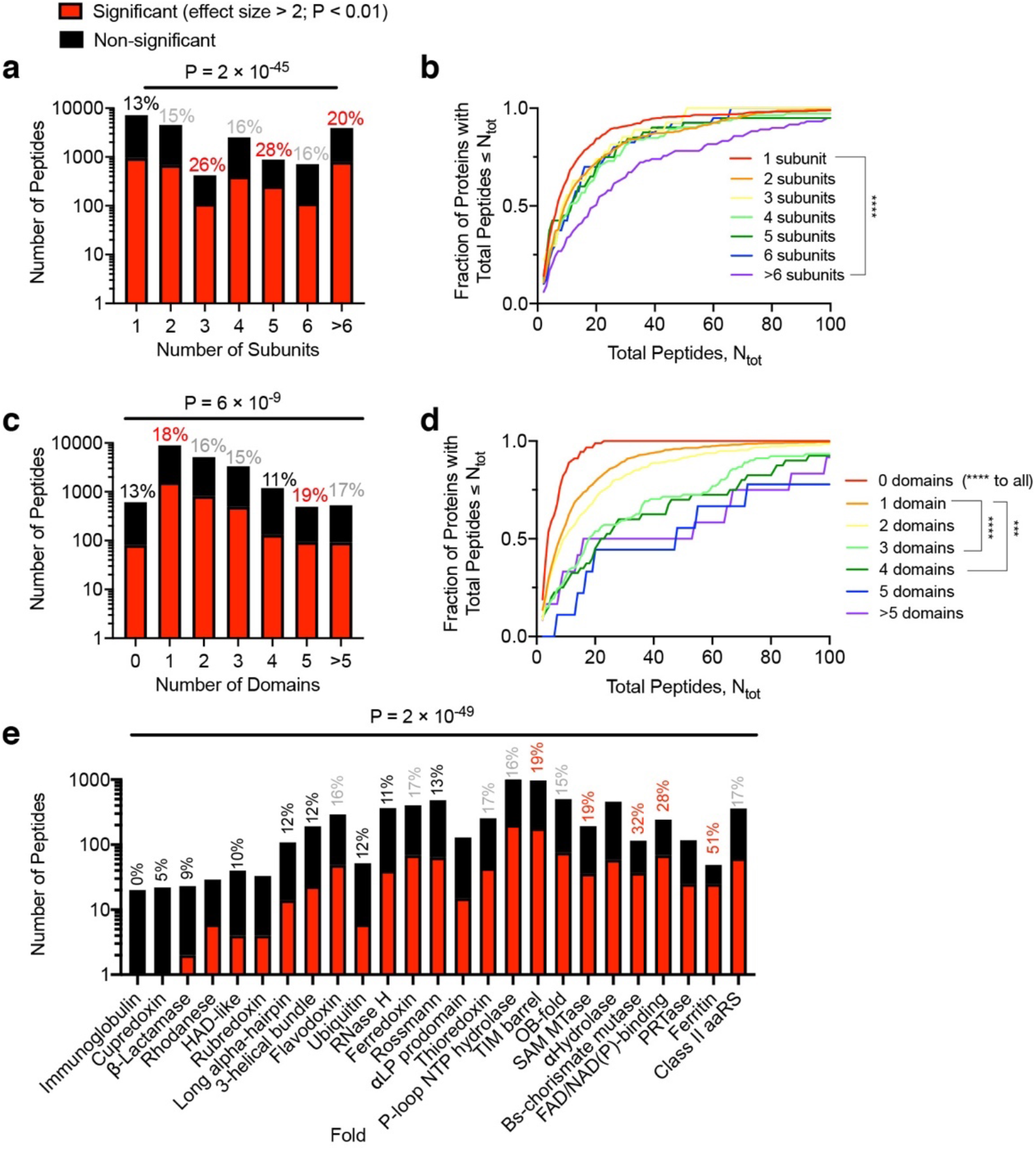
Peptide-Level Analyses to Account for the “*N*_tot_-bias.” The major limitation of the analyses presented in Fig. 3**d,g,i** is that a protein is more likely to be called non-refoldable if we identify and quantify more peptides associated with that protein (see Supplementary Text 3). We call this the *N*_tot_-bias (*N*_tot_ is the total number of peptides quantified for a particular protein). **a,c,e**, Peptide-level analyses in which the total number of significant peptides (red), the total number of non-significant peptides (black), and the fraction of peptides that are significant are given for each classification, *regardless of which protein they came from*. Percentages in black represent a lower significance rate than average, in gray represent a significance rate comparable to average, and in red represent a higher significance rate than average. **b,d**, Cumulative frequency distributions showing the fraction of proteins with *N*_tot_ total peptides quantified or fewer for each classification. This analysis assesses whether proteins of different classifications have significantly different distributions in the total number of peptides per protein, *N*_tot_. **a**, Proteins part of complexes with different numbers of subunits generate significant peptides at different frequencies. Monomeric proteins produce significant peptides at significantly lower frequencies, and proteins part of 3-, 5-, and >6-subunit complexes produce significant peptides at significantly higher frequencies than average (P = 2×10^−45^ by chi-square test), mirroring the protein-level analysis (Fig. 3**d**). **b**, Proteins associated with complexes of different numbers of subunits do not produce different numbers of peptides. By the Kruskal-Wallis test, the distributions are not significantly different from one another, except between the set of monomeric proteins and the set of proteins part of complexes with >6 subunits. This effect is dominated by ribosomal proteins. **c**, Single-domain and 5-domain proteins produce significant peptides at a significantly higher frequencies than average (P = 6×10^−9^ by chi-square test). This does not follow the same trend as the protein-level analysis. **d**, Proteins with more domains tend to produce more peptides by the Kruskal-Wallis test. This effect may be due to the fact that proteins with more domains have more regions that can be targeted by proteinase K. Collectively, these results show that the domain-effect may be potentially biased. **e**, Domains of different fold-types produce significant peptides at significantly different frequencies (P = 2×10^−49^ by chi-square test), mirroring the trends of the domain-level analysis (Fig. 3**i**).

**Extended Data Fig. 6.**
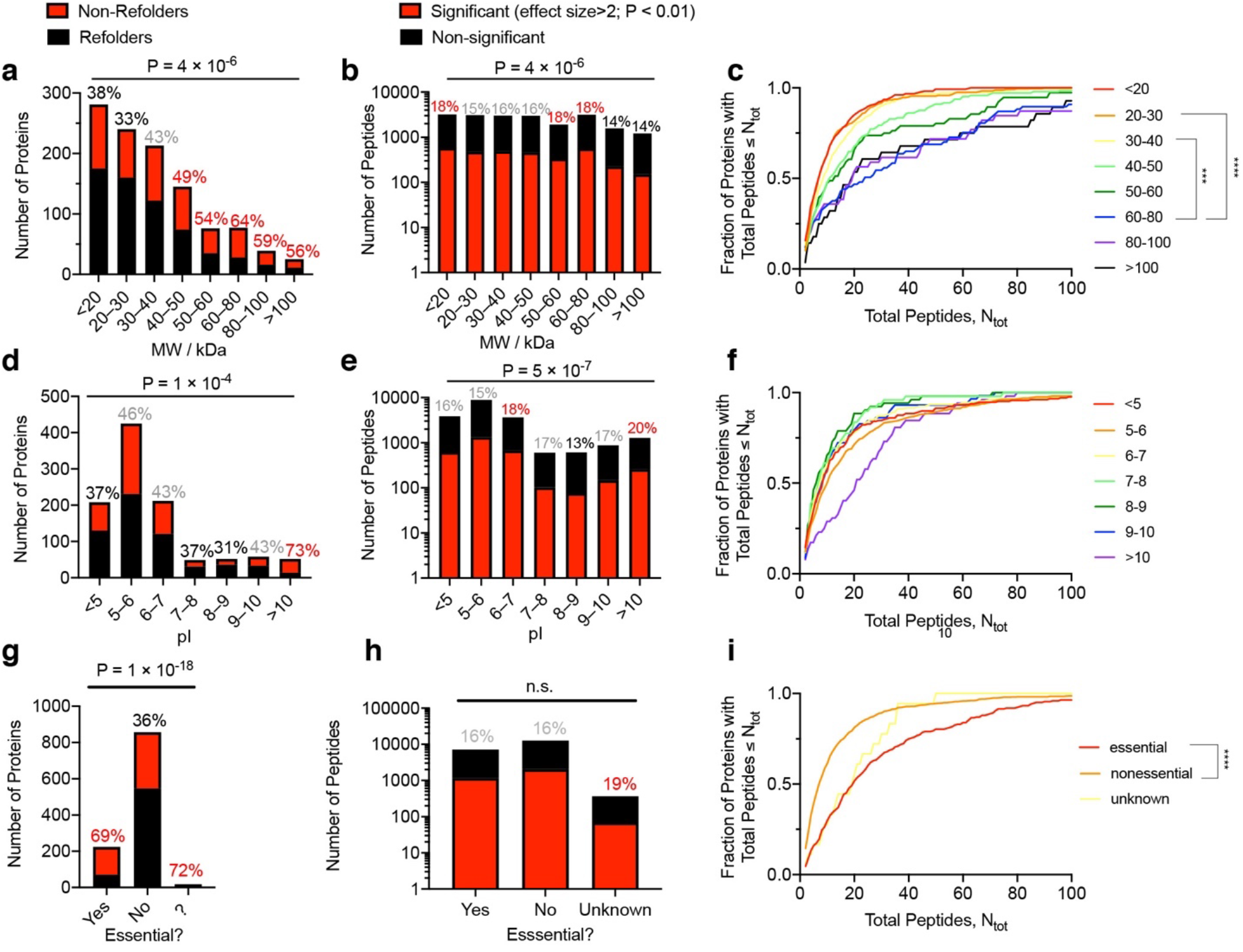
Refoldability of Proteins in terms of Molecular Weight, Isoelectric Point, Essentiality, and Peptide-Level Analyses. **a,d,g**, Refoldability levels at the protein-level, analogous to Fig. 3**d,g,i**. Percentages in black represent a higher refoldability rate than average, in gray represent a comparable refoldability rate to average, and in red represent a higher non-refoldability rate than average. **b,e,h**, Fraction of peptides that are significant for each classification *regardless of which protein they came from*, analogous to Extended Data Fig. 5**a,c,e**. Percentages in black represent a lower significance rate than average, in gray represent a comparable significance rate to average, and in red represent a higher significance rate than average. **c,f,I**, Cumulative frequency distributions showing the fraction of proteins with *N*_tot_ total peptides quantified or fewer for each classification, analogous to Extended Data Fig. 5**b**,**d**. **a**, Proteins of greater molecular weight are significantly more likely to be non-refoldable (P = 4×10^−6^ by Chi-square test). **b**, This effect is partially recapitulated at the peptide-level. **c**, For the most part, proteins of different molecular weights do not produce different numbers of peptides, except for proteins between 60-80 kDa, which produce significantly more than 20-30 kDa and 30-40 kDa proteins by the Kruskall-Wallis test. **d**, Proteins whose isoelectric point (pI) are between 8-9 are significantly more likely to be refoldable, and proteins whose isoelectric point are >10 are significantly more likely to be non-refoldable (P = 1×10^−4^ by Chi-square test). **e**, This effect is recapitulated at the peptide-level, as proteins whose pI are between 8-9 generate significant peptides at a lower frequency, and proteins whose pI are >10 generate significant peptides at a higher frequency than average (P = 5×10^−7^ by Chi-square test). **f**, Proteins of different pI do not produce different numbers of peptides. **g**, Essential proteins are more likely to be non-refoldable (P = 1×10^−18^ by Chi-square test); however, essential proteins do not generate significant peptides at a higher frequency by the Chi-square test, **h**; but they do generate more peptides overall than non-essential proteins by the Kruskall-Wallis test, **i**. Hence, the higher level of non-refoldability of essential proteins is probably biased by their propensity to generate more peptides.

**Extended Data Fig. 7.**
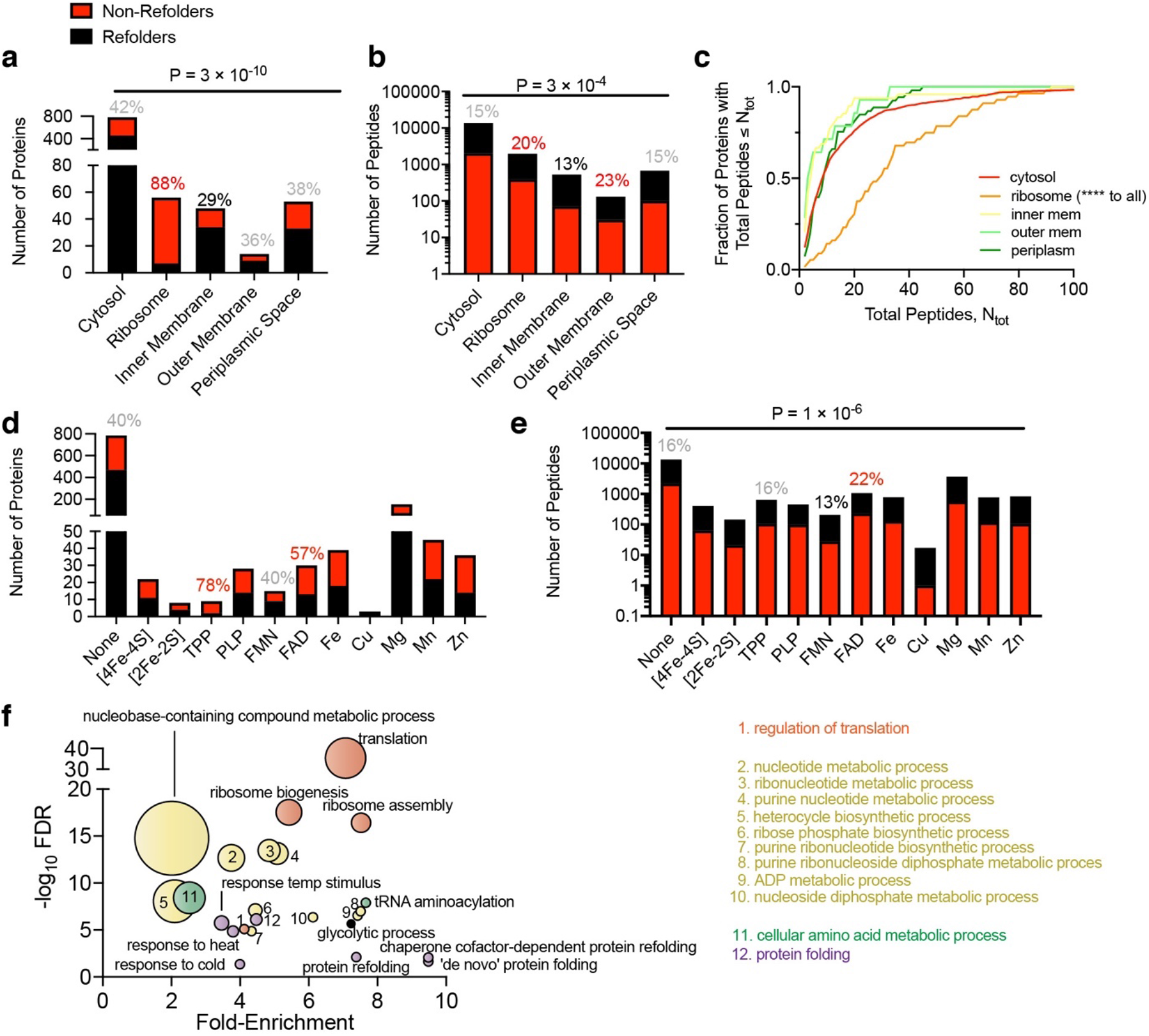
Refoldability of Proteins in terms of Location, Cofactor Presence, and Gene Ontology (GO). **a**, Proteins in the inner membrane are significantly more likely to be refoldable, and proteins that are part of the ribosome are significantly more likely to be non-refoldable than cytosolic or periplasmic proteins (P = 3×10^−10^ by chi-square test). **b**, This effect is recapitulated at the peptide-level, with ribosomal proteins generating significant peptides at a higher rate and inner membrane proteins generating significant peptides at a significantly lower rate than average (P = 3×10^−4^ by chi-square test). **c**, Proteins in different locations do not produce different numbers of peptides by the Kruskall-Wallis test, except for ribosomal proteins (which produce significantly more peptides than others). **d**, Cofactor presence is not associated with major differences in protein refoldability. Proteins with thiamine diphosphate (TPP) or flavin adenine dinucleotide (FAD) cofactors are less refoldable, though this effect is not significant by the chi-square test. **e**, At the peptide level, cofactor differences are statistically significant (P = 1×10^−6^ by chi-square test). TPP-containing proteins do not generate significant proteins at a higher rate, though FAD-containing proteins do. **f**, Gene ontology analysis of non-refolding proteins. Non-refolders are especially enriched amongst proteins involved in the ribosome/translation (red), nucleoside(tide) metabolism – particularly of purines (yellow), aminoacylation (green), and protein folding/stress response (violet).

**Extended Data Fig. 8.**
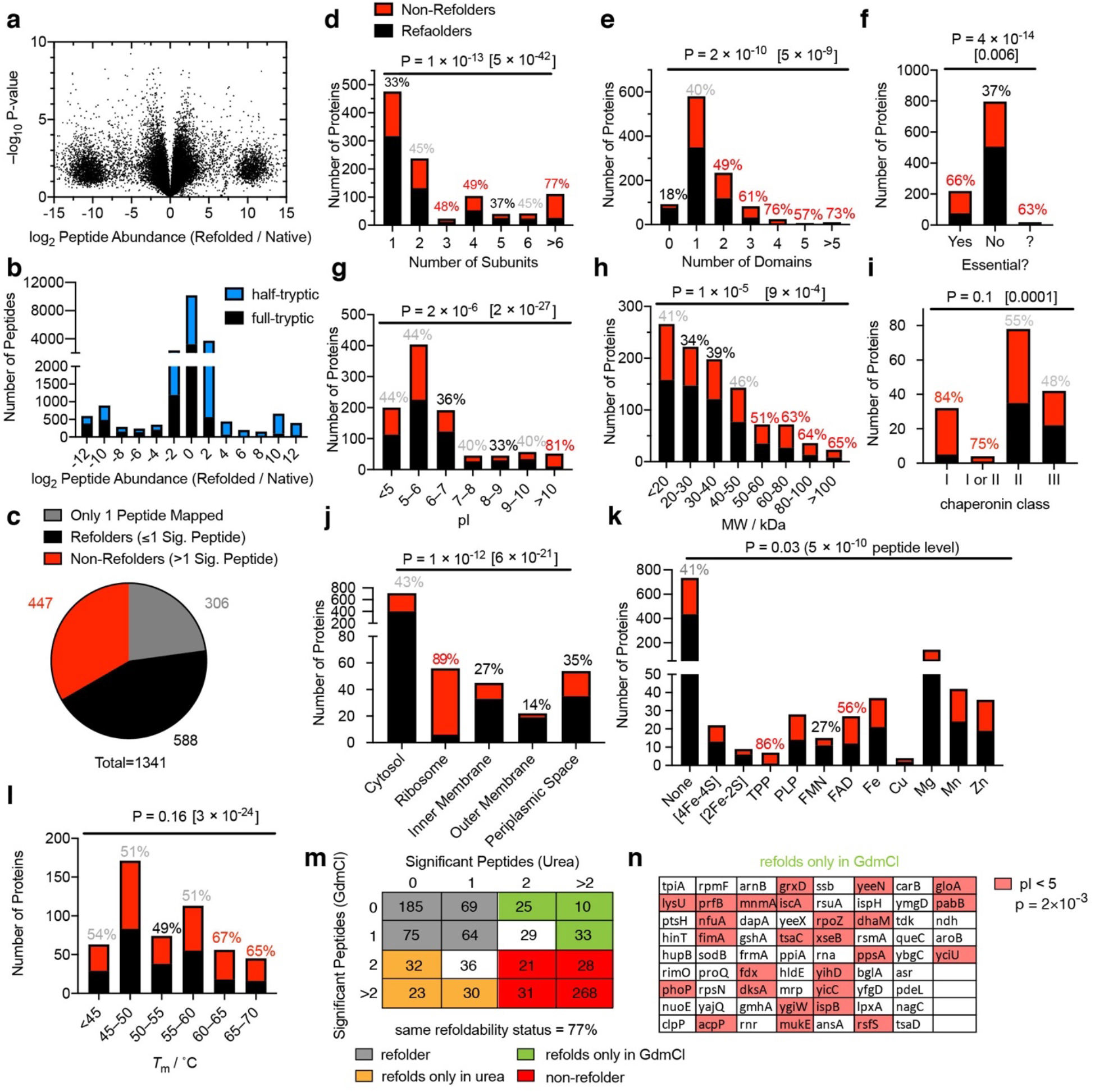
Refoldability of the *E. coli* Proteome from 8 M urea. Figure is analogous to Fig. 3 and Extended Data Figs. 6–7, except for separate experiments in which 8 M urea was used to unfold lysates rather than 6 M GdmCl. The majority of the trends are identical between the two denaturants. **a**, Volcano plot comparing peptide abundances from native and refolded *E. coli* lysate normalized for protein abundance (*n* = 3). Effect sizes reported as ratio of averages, and P-values are based on Welch’s t-test. **b**, Histogram of abundance ratios for half-tryptic and full-tryptic peptides. Half-tryptic peptides are enriched in the refolded proteome. **c**, Overall number of refolding proteins out of 1035 *E. coli* proteins. 306 proteins only furnished one peptide and hence too little data to make an assessment. **d-l**, Bar charts showing number refoldable (black) and non-refoldable (red) proteins for various classifications. Percentages denote percent of proteins that are non-refoldable (red, higher than average; black, lower than average; gray, average). P-values report significance according to the chi-square test; P-values in brackets are conducted at the peptide-level. **d**, Proteins that are part of complexes with multiple subunits are significantly less refoldable than monomeric proteins. **e**, Proteins with more than one domain are significantly less refoldable than proteins with zero or one domain, though this is not recapitulated at the peptide-level. **f**, Essential proteins are significantly less refoldable than non-essential proteins, though this is not recapitulated at the peptide-level. **g**, Proteins with isoelectric point (pI) between 8-9 are the most refoldable, whereas proteins with pI >10 are the least refoldable (P-value = 2×10^−6^ by chi-square test). **h**, Larger proteins are generally less refoldable than smaller proteins. **i**, Class I proteins are the most non-refoldable, whilst class II and class III proteins are evenly split between the categories. **j**, Membrane proteins refold very efficiently out of urea, whereas periplasmic proteins are similar to cytosolic proteins, and ribosome proteins are the least efficient refolders. This is recapitulated at the peptide-level, and is distinct from the GdmCl refolding experiment. **k**, Proteins that harbour TPP and FAD are generally less refoldable; in contrast proteins that harbour FMN are generally more refoldable. This is recapitulated at the peptide-level, and is similar but a stronger effect as compared to the GdmCl refolding experiment. **l**, Highly thermostable proteins are generally non-refoldable, whereas proteins with intermediate thermostability are the most refoldable. Due to low counts, statistical significance is drastically enhanced by conducting analysis at the peptide level. **m**, A total of 1178 proteins were simultaneously identified in the GdmCl and urea experiments, of which 959 had two or more peptides identified for that protein in both the GdmCl and urea experiments, permitting an independent assessment of its refoldability. Table shows the number of proteins that had a given number of significant peptides mapped to it in the GdmCl and urea experiments. 393 were identified as refolders in both experiments and 348 were identified as non-refolders in both experiments. Overall, 77% of proteins had the same refolding status in both experiments. Cells highlighted in green (orange) correspond to proteins that only refold in GdmCl (urea). For this analysis, proteins that were ‘borderline’ (had 1 significant peptide under one condition and 2 significant peptides under the other) were discounted (in white). **n**, List of proteins that only refold from GdmCl. This set of proteins was enriched for those with isoelectric points less than 5 (P = 0.002 by Fisher’s exact test).

**Extended Data Fig. 9.**
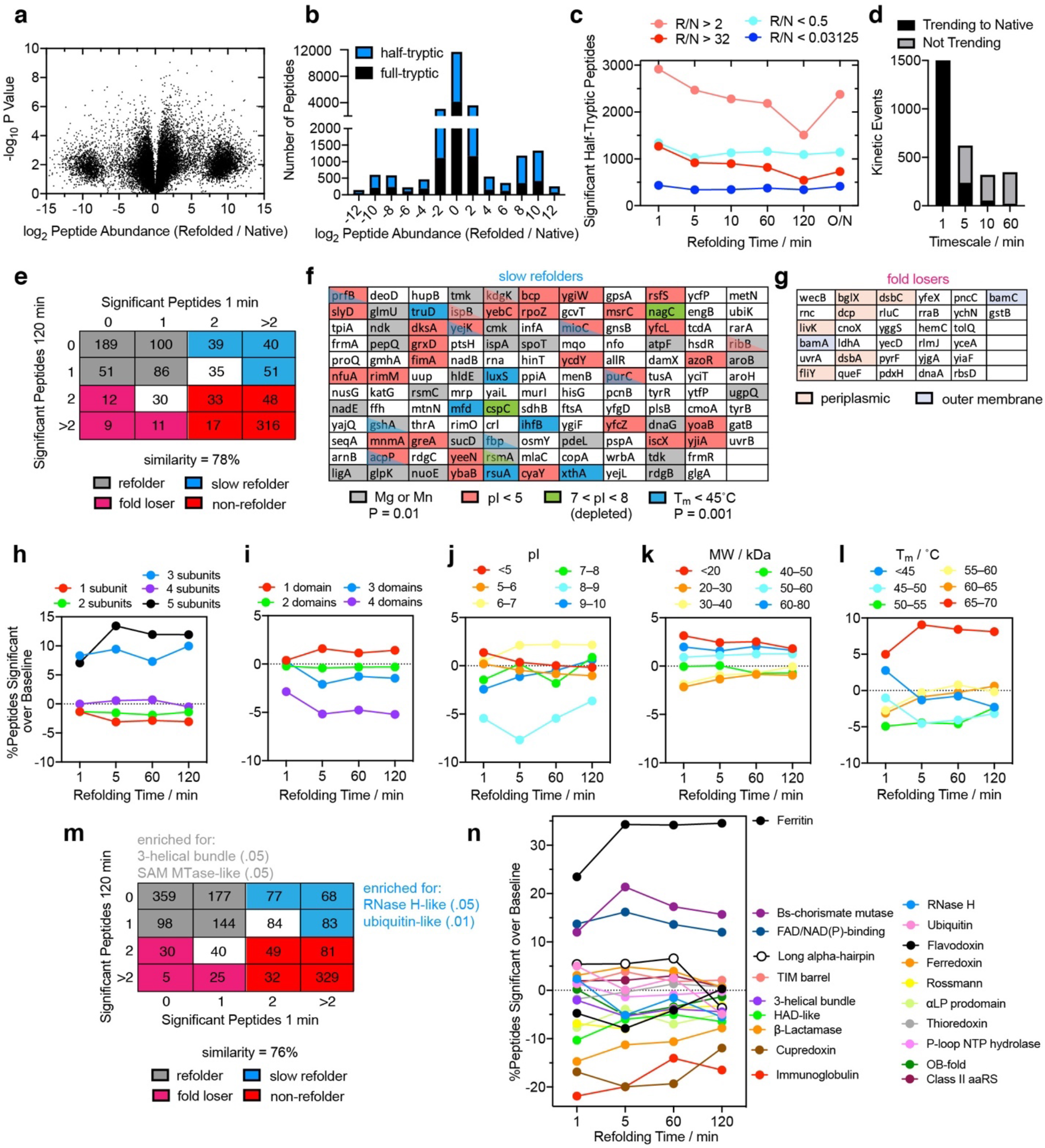
Comparison of Proteome After 1 vs. 120 min of Refolding, and Peptide-level and Domain-level Analysis of Slow Refolding. **a**, Volcano plot comparing peptide abundances ratios from native and refolded *E. coli* lysates normalized for protein abundance (*n* = 3), after 1 min of refolding. Effect sizes reported as ratio of averages, and P-values are based on Welch’s t-test. **b**, Histogram of abundance ratios for half-tryptic and full-tryptic peptides. Half-tryptic peptides are enriched in the 1-min-refolded proteome. Compared to the 120-min-refolded proteome there are significantly more exposed sites (cf. Fig. 3**b**). **c**, Number of significant half-tryptic peptides identified at each time point during kinetic refolding experiments from GdmCl. Peptides are organized into groups based on the size and sign of the effect size (red = site is more exposed in refolded (R); blue = site is more exposed in native (N); darker color = more than 32-fold difference in peptide abundance). **d**, The number of significant half-tryptic peptides that suggest a structural change to become more native-like at various timescales. See Supplementary Text 5. **e**, A total of 1321 proteins were simultaneously identified in the 1-min and 120-min refolding experiments, of which 1067 had two or more peptides identified for that protein at both time points, permitting an independent assessment of its refoldability. Table shows the number of proteins that had a given number of significant peptides mapped to it in the 1 min and 120 min time points. 426 were identified as refolders at both time points and 414 were identified as non-refolders at both time points. Overall, 79% of proteins had the same refolding status at both time points. Cells highlighted in blue correspond to proteins that required more than 1 min to refold (‘slow refolders’). Cells highlighted in magenta correspond to proteins that could refold rapidly but then misfolded afterwards (‘fold losers’). For this analysis, proteins that were ‘borderline’ (had 1 significant peptide at one time point and 2 significant peptides at the other) were discounted (in white). **f**, List of slow refolders. This set of proteins was enriched for those with divalent cation cofactors (P = .01 by Fisher’s exact test) and low thermostability (P = .001 by Fisher’s exact test). **g**, List of fold losers. **h-l**, Variations in refoldability as a function of time for various protein (domain) classifications, with analyses conducted at the peptide level to support the conclusions from Fig. 4**c-f**. To highlight differences, the percent of peptides that are significant for a given classification at a given time is reported as a difference relative to the average frequency. **h**, Proteins in complexes with 2 or 4 subunits are no slower at refolding than monomers. Proteins in complexes with 5 subunits tend to be fold losers. **i**, Proteins with multiple domains are no slower at refolding than single-domain proteins. **j**, Proteins with pI < 5 generate fewer significant peptides as time progresses. **k**, Proteins that weigh <20 kDa generate fewer significant peptides as time progresses. **l**, Proteins with *T*_m_ < 45°C generate fewer significant peptides as time progresses, whereas the more thermostable proteins (*T*_m_ > 60°C) have the opposite trend. **m**, Analogous to **e** but conducted on domains rather than proteins. A total of 2279 domains were simultaneously identified in the 1-min and 120-min refolding experiments, of which 1671 had two or more peptides identified for that protein at both time points, permitting an independent assessment of its refoldability. Table shows the number of domains that had a given number of significant peptides mapped to it in the 1 min and 120 min time points. 778 were identified as refolders at both time points and 491 were identified as non-refolders at both time points. Overall, 76% of domains had the same refolding status at both time points. **n**, Domains of different fold-types evince large differences in their refolding kinetics. Cupredoxin, immunoglobulin, β-lactamse, HAD-like, Rossman, and 3-helical bundle folds refold quickly. Long-alpha hairpins, OB-folds, RNase H-like folds, and TIM-barrels refold slowly. Some folds, such as ferritin-like, FAD/NAD(P) binding domains, and *Bacillus* chorismate mutase-like folds refold poorly at all timescales.

**Extended Data Fig. 10.**
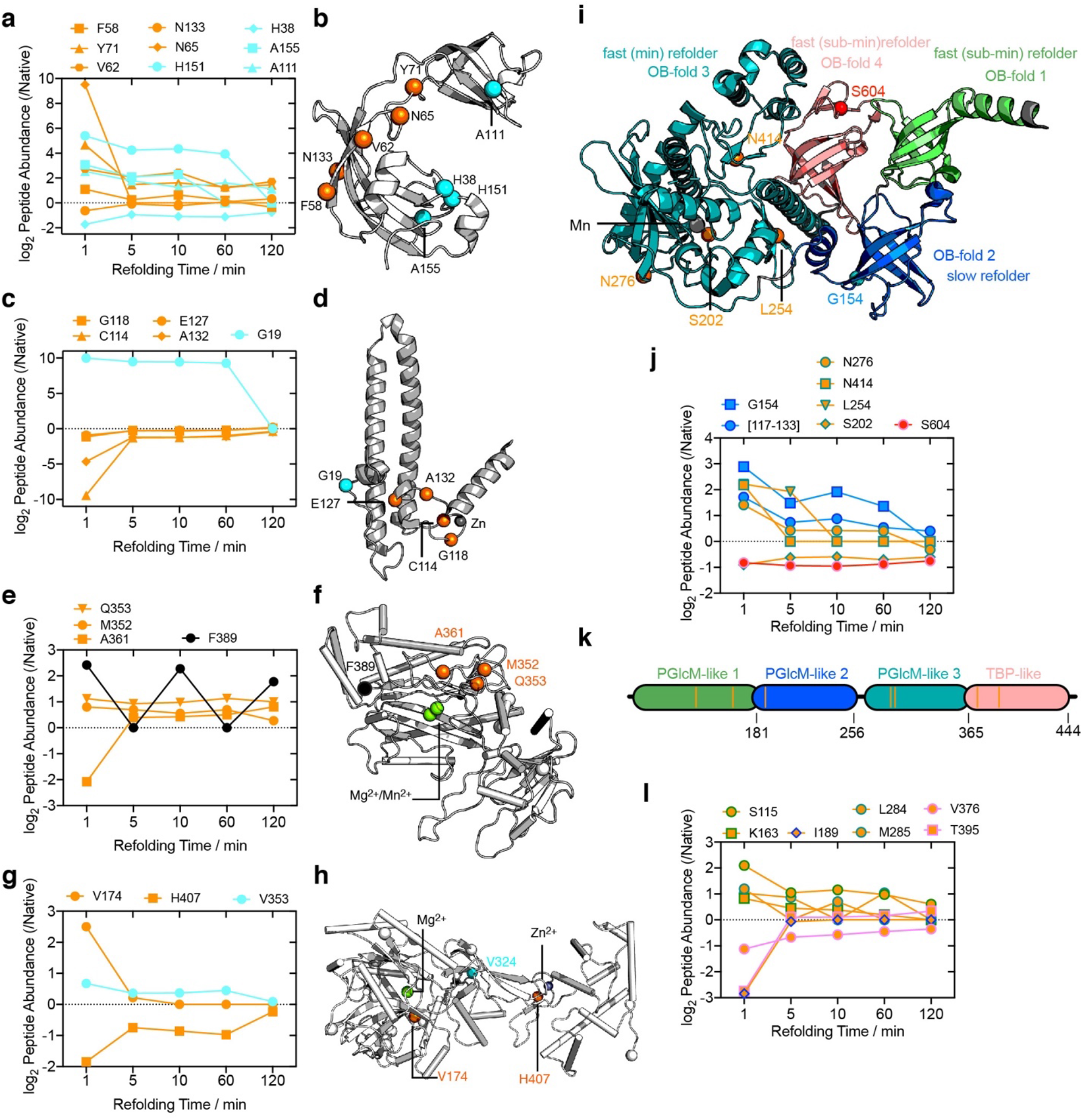
Examples of Minute-Timescale Refolding and Structural Commonalities. Each example displays kinetic traces in panels **a,c,e,g,l** along with structures of the corresponding protein shown in panels **b,d,f,h,i,k**. Orange traces and spheres denote sites which relax to native-like conformation on the 1-min timescale. Cyan traces and spheres denote sites which undergo ‘slow’ processes, relaxing to native-like conformation on slower timescales. **a**, Nine residues of SlyD, a peptidyl-prolyl cis-trans isomerase, whose accessibility to proteinase K change significantly during the min- and h-timescales. **b**, Solution NMR structure of SlyD (PDB: 2K8I). The faster sites cluster around a β-sheet of an FKBP-like domain which is split by an insert-in-flap domain. A series of sites associated with slow refolding are all on one face and may line a putative dimer-interface. **c**, Five residues of DksA, an RNA polymerase-binding transcription factor, whose accessibility to proteinase K change significantly during the min- and h-timescales. **d**, X-ray structure of DksA (PDB: 1TJL). The faster sites cluster around a Zn^2+^ ion and may be due to slow incorporation of the divalent cation. A single slow-evolving site is associated with a kink in a long alpha-hairpin domain. **e**, Four residues of PepQ, an Xaa-Pro dipeptidase, whose accessibility to proteinase K change significantly during the min-timescale. **f**, X-ray structure of PepQ (PDB: 4QR8). The orange sites cluster around a cavity that binds two divalent cations. **g**, Three residues of LigA, *E. coli*’s DNA ligase, whose accessibility to proteinase K change significantly during the min- and h-timescales. **h**, X-ray structure of LigA (PDB: 5TT5). The min-timescale relaxation events are both near divalent cations. **i**, X-ray structure of RNase II (Rnb, PDB: 2ID0), a complex protein with four domains, all of the OB-fold type. **j**, Seven residues whose accessibility to proteinase K change significantly during the min- and h-timescales. The rims of the symbols denote which domain they correspond to. The first domain (green) has no significant peptides at any time point and is presumed a sub-min refolder. The second domain (marine) has one significant site that relaxes slowly. The third domain (teal), which coordinates a Mn^2+^ ion, has four sites which all relax on the min timescale. **k**, Domain structure of phosphoglucosamine mutase (GlmM), for which structures are not available. It consists of three tandem phosphoglucomutase domains, followed by a TBP-like domain. **l**, Seven residues whose accessibility to proteinase K change significantly during the min- and h-timescales. All domains refold on the same timescale. Note also that OB-folds (**i**), long alpha-hairpins (**d**), proteins with low pI (**b,d**), proteins that harbor divalent cations (**d,f,h,i**), and proteins with lower thermostability (**f,h,i**) are all enriched amongst slow-refolders.

