## Supplementary Notes for "Non-Refoldability is Pervasive Across the *E. coli* Proteome"

**Table of Contents**

I. Supplementary Notes

1. Description of Filtering Algorithm for LFQ Data
2. The “$N_{tot}$-bias”
3. A Note on Membrane Proteins
4. Description of Analysis for Refolding Kinetics

II. Supplementary Data Guide

1. K12_GdmCl_summary16Protein.xlsx
2. K12_GdmCl_summary16Domain.xlsx
3. K12_Urea_summary16Protein.xlsx
4. K12_Urea_summary16Domain.xlsx
5. K12_GdmCl_R1-120min.xlsx
6. K12_GdmCl-Urea.xlsx

### Supplementary Notes

#### 1. Description of LFQ Filtering Algorithm

As described in *Materials and Methods,* LFQ analysis comprising of 3 replicates of a refolded sample and 3 replicates of a native sample was conducted in ProteomeDiscoverer (PD) 2.4. For proteomic studies, a separate LFQ analysis comprising of 3 replicates of a refolded control sample and 3 replicates of a native control sample was conducted in PD 2.4. From each .pdResult file, we generated two Excel outputs: a three-tiered output of the results in the hierarchy of Protein>Peptide Group>Consensus Feature (referred to as the PD output file), and a separate file that listed all Consensus Features (referred to as the consensus feature file).

The script called Analyzer_v16_pd24.py (available on GitHub) accepts as inputs the PD output file for the control experiment and the LiP experiment, and the consensus feature file for the LiP experiment.

We use PD's in-built algorithms to infer protein abundance differences based on the available peptide group data for the control experiment (Extended Data Fig. 2e,g). If a protein abundance difference is greater than 2-fold and the P-value calculated by PD is less than 0.01, this was considered to be a significant protein abundance difference and is used as a normalization constant for all peptides that map to that protein in the analysis of the LiP experiment. If either of those thresholds are not met (or if no quantification data is available for said protein, or if that protein was not identified in the control experiment), then no normalization is conducted on that protein.

The main role of the analyzer is to convert the raw extracted ion chromatogram (XIC) peak intensities derived from PD's Minora feature mapper and convert them into an abundance difference to be used for our downstream analysis. Because our analysis relies explicitly on quantification at the peptide level (rather than the protein level) we opted to perform this with in-house scripts. Furthermore, our experiments are characterized by two additional complications not present for most proteomic applications. Firstly, our native and refolded samples – at the peptide level – are very different from each other because of the large number of proteinase K sites that are present only in the native or refolded samples. Hence, retention times can be very different for a given peptide between the two sample-types due to chromatographic matrix effects. Secondly, the fragments that are characteristic of a distinct structure will be absent (or at very low abundance) in either the refolded or native sample, and general LFQ algorithms deal with missing data in a manner that we found unsatisfactory, as discussed in the following.

The hierarchy of the PD output file provides for each protein, a set of peptide groups that maps to that protein; and for each peptide group, a set of consensus features that map to that peptide group. A consensus feature refers to a set of MS1 XICs that were feature-mapped together into a single grouping, *regardless of whether it was successfully assigned to a particular peptide sequence based on MS2*. Hence, each consensus feature consists of a retention time window, a precursor m/z, a charge state, a number of peptide-spectrum matches (PSMs), and a set of intensities for each of the 6 (or more) runs being compared in the LFQ.

For each consensus feature, we assessed whether it should be considered based on the following criteria. If it had zero missing values, it was kept without conditions. If it had one missing value, it was kept though the missing value was discarded (not filled with zero). If it had three non-zero values corresponding to the refolded (or native) replicates and three missing values corresponding to the native (or refolded) replicates, we would only consider it if it was associated with at least 2 or more PSMs. This requirement helps remove cases where the consensus feature may not be correctly assigned to the right peptide group. Next, for each of these situations we searched the consensus feature file (which contained all consensus features, including unassigned ones) for the existence of another consensus feature whose m/z was within 10 ppm of the current consensus feature under consideration and whose retention time was within 5 minutes of the current consensus feature under consideration. If any such consensus feature existed, its set of extracted ion intensities was added to that of the consensus feature under consideration. If not, and there did not exist anywhere in the data any set of intensities that could ‘fill’ the missing values, then we filled them with a default value of 1000 (the approximate detection limit of the instrument, as based on the lowest ion intensities we found in the consensus feature file).

In effect, this procedure 'second-guesses' the software's native feature mapper as extensive manual investigation of our data revealed that in some instances where a peptide appeared to have a very large (or small) abundance ratio, it was because the 'missing' ion intensities existed in the data but were not assigned to the same feature because of a very long retention time shift. This effect was particularly taxing for our experiments because of the large differences in the peptide profile present in the two sample-types we seek to compare. Moreover, application of this procedure treats missing data in a conservative way, and is designed to treat refoldability (which would give rise to abundance ratios close to 1) as a 'global null hypothesis.' This filtering procedure discards (or alters) a significant number of data points with very large (or small) abundance ratios, though leaves behind a subset that are highly reproducible (Extended Data Fig. 3). Overall volcano plots shown in Fig. 3a and Extended Data Figs. 8a and 9a do ***not*** apply this procedure to consensus features with three missing values in order to give an overall picture of how frequent these extreme abundance ratios are. All downstream analyses about refoldability used the filtered data.

After this procedure of discarding (or editing) a consensus feature, we next calculated the ratio associated with that consensus feature (average of the refolded extracted ion intensities divided by the average of the native extracted ion intensities), and a P-value according to the t-test with Welch's correction for non-equal population variance.

Some peptide groups were associated with more than one consensus feature. This occurs frequently if multiple charge states associated with that peptide are detected. It can also happen if the peptide has a propensity to undergo stochastic methionine oxidation. We considered all consensus features for a given peptide group together, including those arising from distinct methionine oxidation levels and different charge states.

If the ratios associated with the various consensus features did not agree in sign (e.g., in the 2+ charge state, the peptide was more abundant in native, but in the 3+ charge state with a methionine oxidation, it was more abundant in refolded), then we assigned the peptide group a ratio of unity (i.e., the data were inconsistent and therefore not used to test against the null hypothesis). If all the ratios agreed in sign, then we took the median of the available ratios as the overall ratio for that peptide group. Moreover, to determine the P-value associated with that ratio we used Fisher's method to combine the P-values associated with the different consensus features. We did not adjust this P-value for multiple hypothesis testing because each set of extracted ion intensities is only used to bear on *one* hypothesis: whether the peptide in question implies identical structure at a given location between the refolded and native forms of a protein, or distinct.

Analyzer compiles each sequenced peptide, along with its associated metadata (see Methods), the identity of the peptide as tryptic or half-tryptic (and if so, the location of the proteinase K cleavage site), abundance ratio, normalized abundance ratio, and P-value and outputs it into a _out.txt file.

Analyzer further performs a protein(domain)-level assessment, whereby it counts the total number of peptides associated with a protein (domain), and the total number that are deemed significant (more than 2-fold abundance difference between native and refolded, P-value less than 0.01). These data are compiled into a _summary16Protein.txt file and a _summary16Domain.txt file (see Supplementary Data). We considered a protein (domain) to be non-refoldable if it had 2 or more significant peptides. We moreover only considered proteins (domains) for the analysis overall if there were 2 or more peptides in total mapped to it. Importantly, the primary claims of the analysis are not sensitive to any of these cut-offs (see Supplementary Data).

**2. The "$N_{tot}$ bias"**

The primary cautionary note one must bear in mind in interpreting our results is the $N_{tot}$ bias. We define $N_{tot}$ as the total number of peptides identified and quantified for a given protein (or domain). Stated simply, because we define a protein (domain) to be non-refoldable if it has two or more peptides that possess significant abundance differences between native and refolded samples ($N_{sig} > 1$), a protein (domain) will be more likely to be judged non-refoldable simply if we quantify more peptides for it. We note that our control studies on SNase and *Tt*RNase H (Fig. 2, Extended Data Fig. 1) indicate that it is, in principle, possible to identify 100-200 unique peptides for a protein, all of which are non-significant. In practice, in our primary experiment (Supplementary Data), the refoldable protein we found with the highest $N_{tot}$ was PrfB (43 peptides), whereas non-refoldable proteins had $N_{tot}$ as high as 254. Hence, we found it prudent to be mindful of this potential source of bias.

We devised two ways to mitigate this bias (see also legend to Extended Data Fig. 5). On one hand, we considered grouping together all significant and non-significant peptides that correspond to a given protein classification (e.g., monomers) without regard to which protein they came from. We refer to this as a peptide-level analysis. The advantage of this approach is that all bias is removed associated with $N_{tot}$ ; moreover, it obviates the need to define a 'minimum' number of significant peptides a protein requires to be labeled non-refoldable. Secondly, we also note that as long as proteins in different classifications that are being compared to each other do not have $N_{tot}$'s that are significantly differently distributed, then we expect the bias to not affect that particular comparison.

The effects reported in this study typically follow one of three courses. If the classifications are confounded by large differences in $N_{tot}$, then the peptide-level analysis gives less significant chi-square P-values; we call these "probably biased." For classifications that do not have large differences in $N_{tot}$, the peptide-level analysis provides enhanced levels of significance; we call these "robust." Finally, for analyses in which the chi-square P-value is roughly the same at the protein and peptide level, we call these "possibly biased."

The differences in non-refoldability of multimers compared to monomers is *robust*, because all the trends at the protein level (Fig. 3d) are recapitulated at the peptide level (Extended Data Fig. 5a) – i.e., monomeric proteins are more refoldable as individual proteins *and* generate significant peptides at a significantly lower rate as a class. The further observation that different subunit classifications have similarly distributed $N_{tot}$'s is reasonable because, for example dimeric proteins would not be expected to be larger, more abundant, or more accessible to proteinase K, on average, than monomeric proteins (Extended Data Fig. 5b) [the significant difference for the >6 subunit classification is affected by the large number of ribosomal proteins].

Another very robust effect is the relationship between fold-type and the refoldability of individual domain. The chi-square P-value at the domain level is modest due to lower counts (Fig. 3i), but is substantially more significant at the peptide-level (Extended Data Fig. 5e).

A more complex relationship is the differences in non-refoldability associated with 'number of domains.' Understandably, a protein with more domains will have more potential proteinase K cut sites, and therefore a higher $N_{tot}$ (Extended Data Fig. 5d). This bias makes itself further known because proteins with higher domain counts do not generate significant peptides at higher rates than single-domain proteins (Extended Data Fig. 5c). Hence, the finding that proteins with more domains are less refoldable is potentially biased.

We found more modest effects linking isoelectric point (pI) and cofactor participation with refoldability (Extended Data Figs. 6d, 7d), although both of these were recapitulated and amplified at the peptide-level (Extended Data Fig. 6e, 7e), so both of these effects are robust. The same was also true for linkages between refoldability and $T_m$ and chaperonin class (see Supplemetary Data).

Essentiality shows a strong relationship with non-refoldability at the protein-level, but hardly any at the peptide level (Extended Data Fig. 6g-h). This can be explained by the fact that essential proteins, generally being larger and more abundant than non-essential proteins, generate more peptides overall (Extended Data Fig. 6i). Hence this result is probably biased.

Finally, the data show unexpected trends between refoldability and molecular weight. Unsurprisingly, the most massive proteins tend toward non-refoldability (Extended Data Fig. 6a), though at the peptide-level these proteins actually generate significant peptides at a lower rate (Extended Data Fig. 6b), so this can be attributed to $N_{tot}$-bias. On the other hand, proteins between 50-60 kDa and 60-80 kDa are more non-refoldable *and* generate significant peptides at a higher rate, so this difference may be real. More unexpectedly, the lightest proteins (<20 kDa) were also more non-refoldable than average-sized proteins *and* generate significant peptides at a higher rate than other weight ranges. We do not currently have an explanation for this observation.

#### 3. A Note on Membrane Proteins

One of the more puzzling results we encountered was the finding that inner membrane proteins are more refoldable than cytosolic proteins (Extended Data Fig. 7a-c), and this effect was even more significant in separate experiments in which proteins were refolded out of urea (Extended Data Fig. 8j). To understand this, we must first consider that cell lysis in our studies is carried out under native conditions by freezer grinding. Consequentially, most membrane proteins are removed from our samples during the clarification of debris from the lysate. However, we hypothesize that freezer grinding can, with some efficiency, liberate soluble portions of membrane proteins, which are then retained in the soluble fraction and subjected to our experimental procedure.

Evidence for this claim comes from the observation that the "membrane proteins" for which we observe the most peptides are more properly considered cytosolic or periplasmic proteins in and of themselves; however, they are part of molecular complexes that are anchored to the membrane. For instance, we observe many peptides for AtpA, AtpC, and AtpD – all components of the cytoplasmic $F_1$ module, and nearly none for AtpE, AtpF, and AtpB – all components of the membrane-bound $F_O$ module. In another example from metabolism, we identify all subunits of the "peripheral arm" of Complex I (NuoBCEFGI) but none of the subunits in the membrane (NuoAHJKLMN). We also observe many peptides for EptC, a membrane protein with a sizable periplasmic domain, but virtually none for SecYEG except for termini, a membrane protein with very little soluble projections (see Supplementary Data).

Hence, our statements about refoldability of inner membrane proteins are best construed as referring to regions of said proteins that reside in aqueous compartments. In this light, the fact that these regions are generally reversibly refoldable is less surprising. Because many inner membrane *E. coli* proteins must be unfolded to be translocated, in Nature they would not be able to benefit from the kinetic coordination of co-translational folding as much as cytosolic proteins, which could fold on the ribosome and never have to be unfolded. Hence, from this point of view, it is easier to rationalize why these portions of inner membrane proteins would be largely refoldable.

**4. Description of Analysis of Refolding Kinetics**

To probe the refolding kinetics of individual proteins, we merged the _out.txt files associated with each time point. Each time point was referenced against a common native-LiP (NL) sample and normalized against a common set of normalization factors obtained from a separate control study (without proteinase K treatment). For time points in which a given peptide was not identified or quantified – resulting in its absence in the corresponding _out.txt file – the peptide was assigned default values of ratio = 1, P-value = 1. As an aside, we adapted analyzer_v16_pd24.py to be able to process LFQ experiments with arbitrarily large number of channels. We found however that more consistent and reliable results were found by performing a series of 3-vs.-3 LFQs and merging the results together.

We next searched for peptides indicative of structural changes that could be kinetically resolved on the min- and h-timescale of this experiment. In the interest of obtaining high-resolution structurally-meaningful results, we only considered half-tryptic peptides pointing to a single proteinase K cut-site. We next checked whether a given peptide was statistically significantly different from native at 1 min (P-value < 0.01). If it was, but it was no longer statistically significantly different at 5 min, then the peptide was judged to imply a kinetic event at the 1-min timescale (Extended Data Fig. 9d). We then asked whether at 5 min, was the abundance ratio more similar to native (closer to 1:1). If so, the peptide was classified as 'trending toward native.' If a peptide was statistically significantly different from native at 1 min and 5 min (but not 10 min), then it was judged to imply a kinetic event at the 5-min timescale. Likewise it was considered to trend toward native if at 1 min, 5 min, and 10 min, the ratio got closer to 1:1. This procedure was continued up until 2 h. As shown in Extended Data Fig. 9d, most of the bona fide kinetic events we could identify were on the min-timescale, with comparatively fewer at longer timescales. For the purpose of counting a peptide as a kinetic event, we did not discard peptides which had a ratio of less than 2-fold, as long as the difference was statistically significantly different from native based on Welch's t-test.

**Supplementary Data**

S1. K12_GdmCl_summary16Protein.xlsx

Summary data for all proteins identified and quantified during refolding experiments from GdmCl. The first six tabs corresponds to different refolding times: 1 min, 5 min, 10 min, 60 min, 120 min, and overnight. Each row corresponds to a protein, denoted with its Uniprot accession code (column A) and gene symbol (column B). Columns C–P and S–U correspond to various metadata for each protein. Columns Q and R report the number of significant peptides and the number of total peptides identified for that protein. Columns AF and AG list the number of refolding and non-refolding proteins associated with various classifications. Columns AI, AJ, and AK list the number of significant peptides, non-significant peptides, and total number of peptides identified that correspond to the given classification (without regard to which protein they came from). Column AL lists the fraction of peptides that are significant. Columns AU and AV show the expected number of refolding and non-refolding protein counts for the purpose of a protein-level chi-square test. Columns AX and AY show the expected number of significant and non-significant peptide counts for the purpose of a peptide-level chi-square test. Down to row 175, these counts are based on using the ‘standard’ rule of a non-refolding protein (two or more significant peptides) and of which proteins are considered for analysis (two or more total peptides). Starting at row 177 is an alternative set of analyses in which all proteins are considered (even those with one peptide only). Starting at row 355 is an alternative set of analyses in which nonrefolders are called for any protein with one or more significant peptides. The last two tabs compile the refoldability rates for all time points at the protein and peptide levels.

S2. K12_GdmCl_summary16Domain.xlsx

Summary data for all domains identified and quantified during refolding experiments from GdmCl. The first six tabs corresponds to different refolding times: 1 min, 5 min, 10 min, 60 min, 120 min, and overnight. Each row corresponds to a domain, denoted with the Uniprot accession code for the protein (column A) and the gene symbol + the residue range (column B). Columns C and D correspond to the fold-type for the given domain and the ordinal position of that domain in its protein. Columns F and G report the number of significant peptides and the number of total peptides identified for that domain. Columns AF and AG list the number of refolding and non-refolding domains associated with various classifications. Columns AI, AJ, and AK list the number of significant peptides, non-significant peptides, and total number of peptides identified that correspond to the given classification (without regard to which domain they came from). Column AL lists the fraction of peptides that are significant. Columns AU and AV show the expected number of refolding and non-refolding domains counts for the purpose of a domain-level chi-square test. Columns AX and AY show the expected number of significant and non-significant peptide counts for the purpose of a peptide-level chi-square test. The last two tabs compile the refoldability rates for all time points at the domain and peptide levels.

S3. K12_Urea_summary16Protein.xlsx

As Data S1, except for experiments in which proteins were unfolded in urea.

S4. K12_Urea_summary16Domain.xlsx

As Data S2, except for experiments in which proteins were unfolded in urea.

S5. K12_GdmCl_R1-120min.xlsx

To generate this file, we considered all proteins which were identified in both the 1 min and 120 min timepoints in the GdmCl refolding experiments. Columns B–S consist of metadata for each protein. Columns T and U report the number of significant peptides and the number of total peptides identified for that protein at 1 min. Columns V and W report the number of significant peptides and the number of total peptides identified for that protein at 120 min. Columns X–AA assign a "1" to each protein that is classified as a slow refolder, fold loser, always refolder, or always non-refolder. Columns AJ–AL list the number of always-refolding, slow refolding, and fold-losing proteins associated with various classifications. Columns AP-AR show expected values for the purpose of protein-level chi-square test. The second tab uses a similar formalism but classifies at the level of domains.

S6. K12_GdmCl_Urea.xlsx

To generate this file, we considered all proteins which were identified in both the GdmCl/120 min timepoint and the urea/overnight timepoint. Columns B–S consist of metadata for each protein. Columns T and U report the number of significant peptides and the number of total peptides identified for that protein when refolded out of GdmCl. Columns V and W report the number of significant peptides and the number of total peptides identified for that protein when refolded out of urea. Columns X–AA assign a "1" to each protein that refolds only out of urea, refolds only out of GdmCl, always refolds, or does not refold in either. Columns AJ–AL list the number of proteins associated with various classifications that refolds only out of urea, refolds only out of GdmCl, or always refolds. Columns AP-AR show expected values for the purpose of protein-level chi-square tests. The second tab uses a similar formalism but classifies at the level of domains.
